# TSC/mTORC1-dependent proteostasis in stress-recruited prefrontal ensembles gates flexible avoidance

**DOI:** 10.64898/2026.11.21.768270

**Authors:** Saheed Lawal, Dhananjay Sharma, Olivia Tabaka, Moniza Mujtaba, Prerana Shrestha

## Abstract

Adaptive threat responses require accurate appraisal of danger signals and flexible behavioral control. Anxiety and trauma-related disorders are marked by avoidance that outlasts the circumstances warranting it, yet such states rarely arise from genetic or environmental insult alone. Here we show that the two converge on a defined prefrontal population by releasing the TSC complex’s constitutive brake on mTORC1. Using activity-dependent genetic targeting in mice, we accessed medial prefrontal cortex neurons recruited during the early phase of social isolation - a population projecting predominantly to the lateral septum - and deleted Tsc2 selectively within them. Ensemble-specific Tsc2 haploinsufficiency suppressed autophagic flux without altering bulk translation and produced behavioral avoidance that was rapid and efficient yet no longer under stimulus control, escalating into extra-CS epochs and generalizing to a novel, non-reinforced context. Mutant animals bypassed the phase in which freezing and shuttling compete, and in vivo calcium fiber photometry revealed ensembles locked to the cue but unable to signal trial outcome or track the competing Pavlovian response. Reinstating restraint on mTORC1 through ensemble-restricted Raptor knockdown or systemic Rapamycin restored stimulus control over avoidance, whereas Tsc2 haploinsufficiency in mPFC neurons without social isolation produced no phenotype. These findings identify TSC-restrained, mTORC1-dependent proteostasis within stress-recruited ensembles as a gene-by-environment locus for maladaptive avoidance. More broadly, they suggest that excessive avoidance in anxiety and post-traumatic stress disorders may arise not from genetic risk or adversity alone but from their convergence on mTORC1 signaling cascade.

## Introduction

Survival depends on the capacity to detect environmental threats and mount calibrated behavioral responses. Adaptive fear learning allows organisms to recognize danger signals, mobilize defensive behavior, and update those responses as contingencies change. When this system becomes dysregulated, threat responses generalize inappropriately, avoidance becomes rigid, and fear persists long after danger has passed ^1–3^. Exaggerated threat appraisal, compulsive avoidance, and failure of extinction are defining features of anxiety and trauma-related disorders, the most prevalent class of psychiatric illness worldwide ^4–6^. Yet the mechanisms that transform adaptive threat learning into pathological behavioral states remain incompletely understood - in part because such states rarely arise from genetic or environmental insult alone, but from their convergence.

The medial prefrontal cortex (mPFC) is a likely site of the convergence between genetic vulnerability and environmental impact. It integrates bottom-up signals from the amygdala, hippocampus, and hypothalamus with top-down control to arbitrate defensive responding ^7–9^, and it is among the structures most sensitive to chronic stress: dendritic retraction, altered excitatory-inhibitory balance, and changes in projection neuron activity accompany anxiety, anhedonia, and impaired extinction ^10–13^. Chronic social isolation - prolonged deprivation of conspecific contact - is a well-validated model of psychosocial adversity that induces persistent negative affect, heightened threat reactivity, and maladaptive avoidance across species ^14–16^. Epidemiologically, isolation and loneliness elevate risk for anxiety, depression, and post-traumatic stress disorder, particularly in individuals with pre-existing neurodevelopmental vulnerability ^17,18^. Several studies over the last decade have shown that experience is not encoded uniformly across a region but within sparse, co-activated ensembles that are necessary and sufficient to drive specific behavioral states and are molecularly distinguishable from surrounding neurons ^19–23^. This raises the possibility that stress acts not on the mPFC as a whole, but on a defined population within it - and that vulnerability is conferred by what happens inside those cells.

Mammalian target of rapamycin complex I (mTORC1) signaling is a strong candidate for cellular mechanism conferring such vulnerability. The TSC1/TSC2 complex is a central negative regulator of mTORC1. TSC2 acts as a GTPase-activating protein for the small G protein Rheb; loss of the complex leaves Rheb constitutively GTP-loaded and mTORC1 persistently active ^24–26^. Under normal conditions, mTORC1 integrates growth factor, nutrient, and synaptic activity signals to reciprocally gate the two arms of proteostasis: it promotes translation initiation by phosphorylating 4E-BP1/2 and S6K1 and simultaneously suppresses degradation by phosphorylating ULK1 to block autophagosome initiation and TFEB to sequester it from the nucleus, shutting down the transcriptional program that replenishes lysosomal capacity ^27–30^. mTORC1 thus exerts two coordinated and antagonistic outputs - promoting protein synthesis while suppressing protein turnover - that are regulated in an experience-dependent manner. Loss of Tsc1 or Tsc2 removes the brake and neither arm remains regulated: translation proceeds without gating while autophagic flux is chronically suppressed. The consequence is not simply excess synthesis but loss of turnover, which is particularly consequential in post-mitotic neurons, where autophagy mediates synaptic remodeling, receptor turnover, and spine elimination - requirements for the bidirectional plasticity underlying normal learning ^31–35^. Structural correlates in TSC models, including elevated spine density and impaired pruning, are consistent with impaired degradation in the context of preserved synthesis ^32,36–38^. In humans, germline loss of Tsc1 or Tsc2 causes tuberous sclerosis complex, a multisystem disorder affecting ∼1 in 6,000 individuals that produces not only cortical tubers and subependymal nodules but a broad neuropsychiatric phenotype, including anxiety and emotional dysregulation in a substantial proportion of patients ^39–42^. How dysregulated proteostasis produces these affective symptoms has remained unclear, because prior work has not resolved the neural populations in which proteostasis must be lost for emotional pathology to emerge.

Here we test this directly, using activity-dependent genetic targeting ^19,43,44^ to access mPFC neurons recruited during the early phase of chronic social isolation, and show that haploinsufficiency of Tsc2 restricted to these stress-activated ensembles produces avoidance that escalates beyond the conditions licensing it. In a signaled active avoidance task ^45–47^, mutant animals bypass the competitive phase between freezing and shuttling that normally precedes efficient responding, and their shuttling progressively invades extra-CS epochs in the absence of any threat cue. Tsc2 haploinsufficiency across prefrontal neurons in animals that never experience isolation does not recapitulate this phenotype, establishing that the genetic lesion is consequential only where stress has recruited the cells in which it acts. Within the ensemble, Tsc2 haploinsufficiency suppresses autophagic flux without altering bulk translation and yields somatodendritic hypertrophy; these neurons are driven by the conditioned cue from the first training session and fail to habituate to the shock yet no longer discriminate successful from failed trials or reciprocally track the competing freezing response. Ensemble-targeted knockdown of the obligate mTORC1 component Raptor, or pharmacological inhibition with rapamycin, restores the vigor of avoidance and confines responding to the cue. These findings identify dysregulated mTORC1 signaling within stress-recruited prefrontal ensembles as a causal mechanism for maladaptive avoidance and show how environmental adversity and disrupted proteostasis converge on a defined cellular substrate to reshape defensive behavior.

## Results

### 1. Social isolation truncates response competition during avoidance learning and recruits prefrontal-septal neurons

To determine how social isolation stress (SIS) affects aversive learning and memory, adult wildtype mice were transferred from group housing to single housing starting two weeks prior to behavioral testing (**Fig. 1A**); controls remained group-housed (GH) throughout. We used the signaled active avoidance (SAA) paradigm, in which mice learn to shuttle between compartments to avoid a footshock signaled by a conditioned stimulus (CS) tone (**Fig. 1B**). The first training session (SAA1) began with a single Pavlovian trial to establish the tone-shock contingency, followed by 11 CS presentations^48^. Shuttling within 15 s of CS onset (CS_max_) was scored as a successful avoidance; failure to shuttle during the CS resulted in delivery of the footshock unconditioned stimulus (US). Successful avoidance requires resolving a conflict between two competing defensive strategies: reactive coping, expressed as Pavlovian freezing, and proactive coping, expressed as instrumental shuttling (**Fig. 1C**). Animals must suppress the freezing response to execute the action that prevents the shock. SAA1 was followed by two additional days of training in the same context (SAA1 and SAA2).

**Figure 1.**
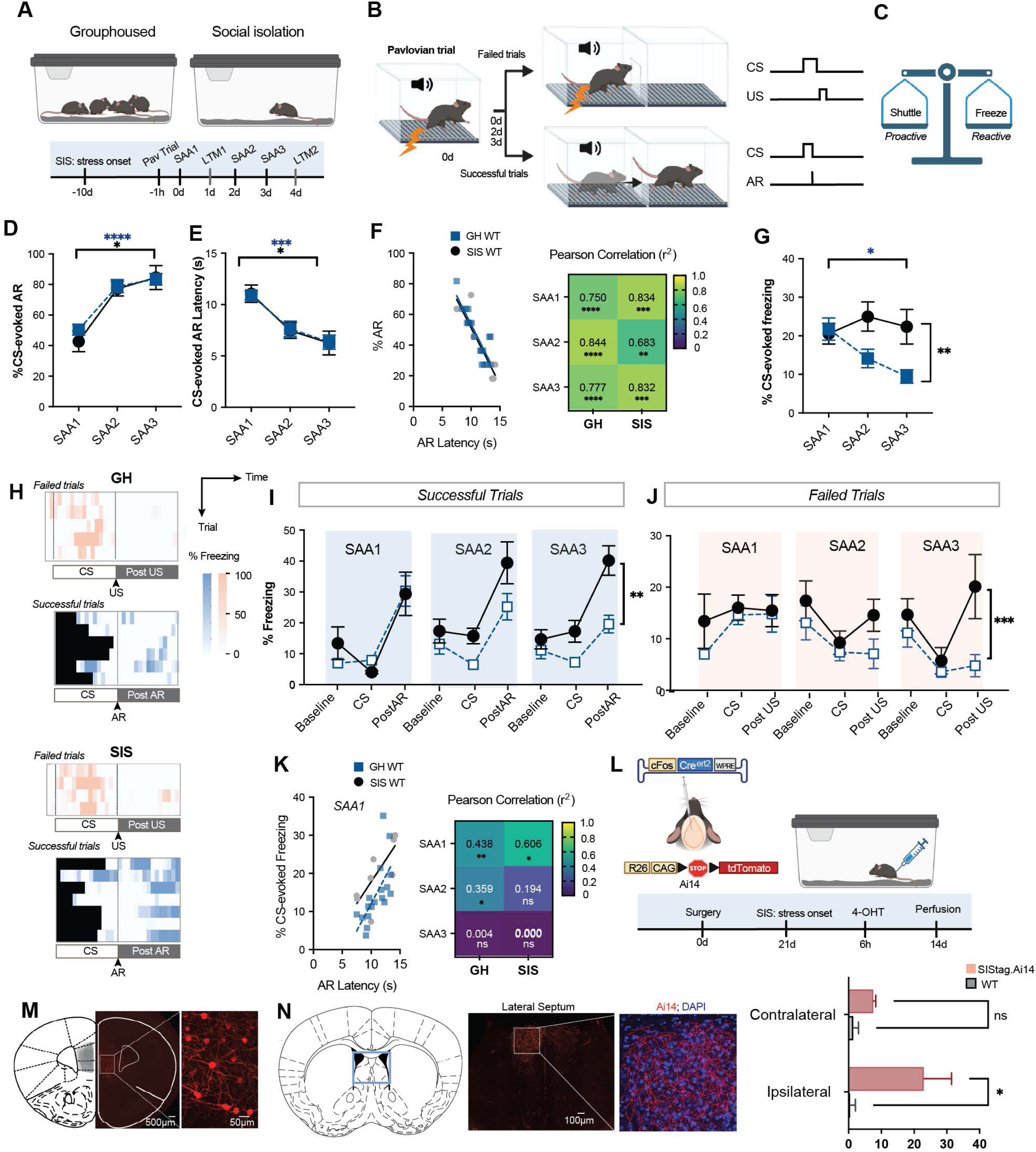
Social isolation truncates response competition during avoidance learning and recruits prefrontal-septal neurons. **A.** Experimental design. Upper, social isolation stress (SIS) paradigm in which mice were singly housed for 10 days before behavioral testing. Lower, timeline of Pavlovian conditioning and signaled active avoidance (SAA). Mice received a Pavlovian conditioning trial and the first SAA training session (SAA1) on day 0, followed by long-term memory test 1 (LTM1) 24 h later, additional reinforced SAA training sessions (SAA2 and SAA3) on days 2 and 3, respectively, and long-term memory test 2 (LTM2) on day 4. **B**. Schematic of the SAA task. Left, training context illustrating successful avoidance trials, in which mice shuttled during the conditioned stimulus (CS), and failed trials, in which the unconditioned stimulus (US) was delivered. Right, task structure showing that the CS terminated upon a successful avoidance response (AR), whereas failure to shuttle resulted in immediate US delivery. **C**. Conceptual model illustrating active avoidance behavior as the outcome of competition between proactive (avoidance) and reactive (freezing) defensive responses. **D**. Percentage of CS-evoked avoidance responses (AR) across SAA1–SAA3. No significant differences were observed between group-housed (GH; blue) and SIS (black) mice. **E**. Latency to initiate the avoidance response during SAA1–SAA3. No significant group differences were detected. **F**. Left, correlation between avoidance performance (%AR) and avoidance latency across SAA1-SAA3, showing strong negative correlations in both GH and SIS mice. Right, summary of Pearson correlation coefficients (r²) and significance values for each training session. **G**. Percentage of CS-evoked freezing during SAA1–SAA3. SIS mice exhibited persistently elevated freezing relative to GH controls. **H**. Representative freezing heatmaps during failed (top) and successful (bottom) trials aligned to the CS and post-US or post-AR periods. **I**. Freezing during successful trials quantified across baseline, CS, and post-AR epochs. SIS mice displayed elevated freezing during the CS epoch relative to GH controls. **J**. Freezing during failed trials quantified across baseline, CS, and post-US epochs. SIS mice exhibited greater freezing, particularly during the post-US epoch of SAA3. **K**. Left, correlation between CS-evoked freezing and avoidance latency across SAA1–SAA3, demonstrating positive correlations in both GH and SIS mice. Right, summary of Pearson correlation coefficients (r²) and significance values for each training session. **L**. Genetic strategy for activity-dependent labeling of the social isolation ensemble using cFos-CreERT2 expression in Ai14 mice. Mice underwent social isolation followed by 4-hydroxytamoxifen administration to permanently label neurons activated during stress. **M**. Representative images of tdTomato-labeled social isolation ensemble neurons in the medial prefrontal cortex (mPFC). Scale bar = 500 μm, 50 μm. **N**. Axonal projections of the labeled stress ensemble to the lateral septum (LS). Left and middle, representative images. Scale bar = 100 μm Right, quantification of ipsilateral and contralateral axon density. **Statistical analyses**: Data are presented as mean ± SEM. Panels D,E,G: Repeated measures two-way ANOVA with Bonferroni post hoc test; Panels F&K: Pearson’s correlation with coefficients (r²) reported; Panels I&J: Mixed-effects analysis with Bonferroni post hoc test; Panel N: Two-way ANOVA with Bonferroni post hoc test. Sample size: **N** = 9 - 22 mice per group (Panels D,E,F,G,I,J,K); **N** = 3 mice per group (Panel N). *p < 0.05, **p < 0.01, ***p < 0.001, ****p < 0.0001; ns, not significant. Full statistical results are provided in **Extended Data Table 1**.

Group-housed (GH) and socially isolated (SIS) wildtype mice acquired CS-evoked avoidance responses at comparable rates, with both groups increasing avoidance across successive training sessions (**Fig. 1D**). Avoidance latency declined in parallel (**Fig. 1E**), indicating that animals not only shuttled more frequently during the CS but did so more rapidly as training progressed; accordingly, percentage avoidance and mean avoidance response (AR) latency were strongly negatively correlated across all SAA sessions (**Fig. 1F**). If shuttling and freezing compete, CS-evoked freezing should decline as avoidance is acquired. This held in group-housed mice, which progressively reduced CS-evoked freezing across sessions. Socially isolated mice, by contrast, maintained high and inflexible freezing throughout all three sessions despite performing avoidance responses at normal rates - a dissociation between the Pavlovian and instrumental responses that is absent in controls (**Fig. 1G**). Resolving freezing by epoch, both group-housed and socially isolated mice froze more in the peri-CS window after an avoidance response (postAR) than at baseline or during the CS, indicating that avoidance was fear-motivated (**Fig. 1H, I**). On failed trials, freezing was minimal at baseline, CS, and post-US epochs in both groups, except for SAA3, when socially isolated mice froze substantially following the US (**Fig. 1H, J**). CS-evoked freezing and AR latency were strongly positively correlated in group-housed mice during SAA1 and SAA2, but in socially isolated mice only during SAA1 (**Fig. 1K**). Because this correlation reflects the cost that freezing imposes on the latency to shuttle, its persistence indexes the period over which the two responses actively compete. Social isolation therefore truncates the competitive phase to a single session, after which avoidance proceeds without being gated by the freezing response it must otherwise overcome - despite freezing remaining high.

We next asked which mPFC neurons are recruited by social isolation itself. Ai14 mice ^49^, which express Cre-dependent tdTomato reporter, received mPFC injections of AAV5-cFos-CreERT2 ^43^, permitting activity-dependent labelling of neurons active within a defined temporal window (**Fig. 1L**). Twenty-one days after surgery, mice were separated from their cagemates and singly housed; 4-hydroxytamoxifen was administered 6 h later, opening a labelling window of approximately 8 h that captured the early phase of isolation. Following 21 days of tdTomato expression, mice were perfused and labelled neurons quantified. This strategy robustly tagged approximately 60 SIS-activated neurons in each hemisphere of the mPFC (**Fig. 1M**). Characterizing their projection targets, we found that the majority projected to the ipsilateral lateral septum (**Fig. 1N**), indicating that prefrontal-septal projection neurons are preferentially recruited during the early isolation period. These results thus identify a prefrontal-septal population engaged at the onset of isolation, coincident with the behavioral signature of freezing decoupled from avoidance.

### 2. Tsc2 haploinsufficiency in stress-activated mPFC neurons suppresses autophagy and produces maladaptive avoidance

To selectively disrupt Tsc2 in mPFC neurons activated by social isolation stress, we employed a conditional heterozygous mouse line carrying a floxed Tsc2 allele ^50^, with wild-type (WT) C57BL/6J mice serving as controls. Both groups received stereotaxic co-injection of AAV5-cFos-CreERT2 and AAV-CAG-DIO-tdTomato into the mPFC (**Fig. 2A**). This dual-virus approach confers tamoxifen-inducible Cre recombinase expression specifically in neurons displaying stress-evoked c-Fos, thereby enabling activity-dependent Tsc2 recombination in Tsc2 floxed mice and permanent fluorescent labeling of the stress-responsive ensemble. In SIStag.TSC mice, tdTomato-positive cells displayed efficient *Tsc2* inactivation, evidenced by a marked reduction in TSC2 protein levels (**Fig. 2B, C**). Morphometric analyses revealed pronounced structural remodeling within the SIS tagged ensemble of mutant mice. tdTomato+ cells in SIStag.TSC mPFC exhibited enlarged soma size (**Fig. 2D**) and increased dendritic arborization compared to WT (**Fig. 2E**). We next asked how heterozygous Tsc2 loss affects the two arms of mTORC1-dependent proteostasis within these cells. To assess nascent protein synthesis, we performed *ex vivo* fluorescent non-canonical amino acid tagging (FUNCAT)^51^, in which the L-methionine analog L-azidohomoalanine is incorporated into elongating polypeptides and visualized by click chemistry. Nascent protein synthesis in SIStag.TSC ensembles was comparable to SIStag.WT (**Fig. 2F**). Autophagic flux, by contrast, was markedly reduced in mutant cells, as indicated by accumulation of p62 (2.1-fold increase) (**Fig. 2G**). Since p62 is an autophagy receptor that is itself degraded upon autophagosome-lysosome fusion, its accumulation reflects impaired autophagic degradation rather than increased synthesis ^52,53^. Heterozygous Tsc2 deletion in stress-recruited mPFC neurons therefore suppresses autophagy while leaving bulk translation intact, dissociating the two arms downstream of mTORC1.

**Figure 2:**
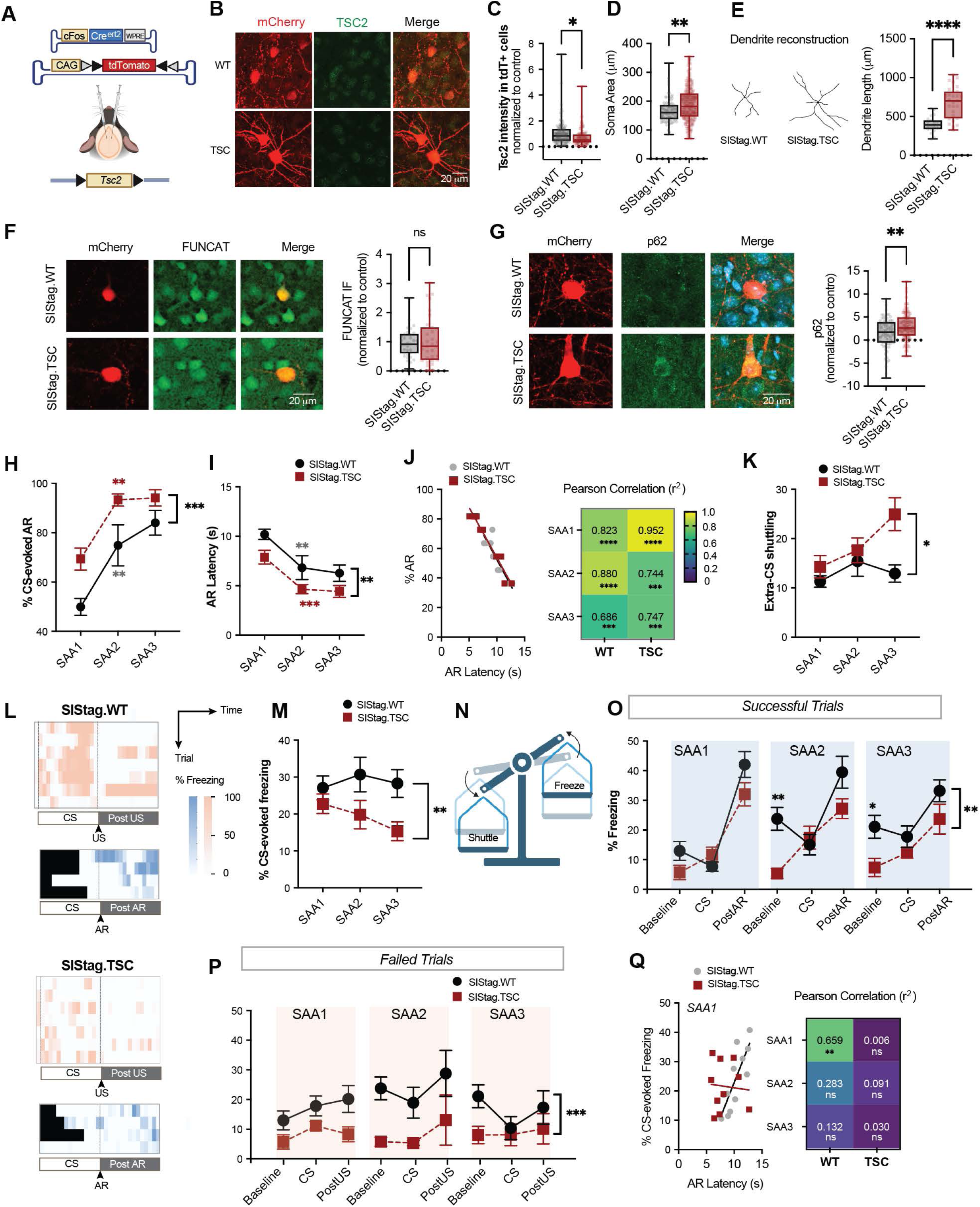
Tsc2 heterozygous loss in stress-activated mPFC neurons suppresses autophagy and produces maladaptive avoidance. **A.** Genetic strategy for activity-dependent labeling of the social isolation ensemble and conditional haploinsufficiency of *Tsc2* by co-injecting AAV5-cFos-CreERT2 and AAV-CAG-DIO-tdTomato into the mPFC of *Tsc2* floxed (SIStag.TSC) and wild-type (SIStag.WT) mice. **B.** Representative images of TSC2 immunofluorescence in SIStag.WT and SIStag.TSC neurons. Scale bar = 50 μm. **C.** Quantification of TSC2 fluorescence intensity demonstrating a significant reduction in SIStag.TSC (red) relative to SIStag.WT (black). **D.** Quantification of neuronal soma area showing enlarged cell size in SIStag.TSC compared with SIStag.WT neurons. **E.** Left, representative dendritic reconstructions of SIStag.WT and SIStag.TSC neurons. Right, quantification of total dendritic length demonstrating increased dendritic arborization in SIStag.TSC neurons. **F.** Left, fluorescence non-canonical amino acid tagging (FUNCAT) assay for nascent protein synthesis. Scale bar = 50 μm. Right, quantification showing no significant difference in bulk protein synthesis between SIStag.WT and SIStag.TSC neurons. **G.** Left, representative images of p62 immunofluorescence in SIStag.WT and SIStag.TSC neurons. Scale bar = 50 μm. Right, quantification of p62 fluorescence intensity demonstrating a significant reduction in SIStag.TSC relative to SIStag.WT neurons. **H.** Percentage of CS-evoked AR across signaled active avoidance (SAA) sessions (SAA1-SAA3). SIStag.TSC mice exhibited increased avoidance responses relative to SIStag.WT controls. **I.** Latency to initiate the avoidance response during SAA1-SAA3. SIStag.TSC mice displayed shorter avoidance latencies than SIStag.WT mice across training. **J.** Left, correlation between avoidance performance (% AR) and AR latency across SAA1-SAA3, demonstrating strong negative correlations in both SIStag.WT and SIStag.TSC mice. Right, summary of Pearson correlation coefficients (r²) and significance values for each training session. **K.** Extra-CS shuttling during SAA training. SIStag.TSC mice exhibited increased extra-CS shuttling during SAA3 relative to SIStag.WT controls. **L.** Representative freezing heatmaps during failed (top) and successful (bottom) trials aligned to the CS and post-unconditioned stimulus (US) or post-avoidance response (AR) periods. **M.** Percentage of CS-evoked freezing across SAA1-SAA3. SIStag.TSC mice exhibited reduced freezing relative to SIStag.WT controls. **N.** Conceptual model illustrating that *Tsc2* haploinsufficiency in stress-activated mPFC neurons biases defensive behavior toward proactive avoidance at the expense of reactive freezing. **O.** Freezing during successful trials quantified across baseline, CS, and post-AR epochs. SIStag.TSC mice exhibited reduced freezing during the baseline epoch of SAA2 and across all epochs during SAA3 relative to SIStag.WT mice. **P.** Freezing during failed trials quantified across baseline, CS, and post-US epochs. SIStag.TSC mice exhibited reduced freezing across all epochs compared with SIStag.WT controls. **Q.** Left, correlation between CS-evoked freezing and avoidance latency during SAA1, revealing a positive correlation in SIStag.WT mice but not in SIStag.TSC mice. Right, summary of Pearson correlation coefficients (r²) and significance values for each training session. **Statistical analyses**: Data are presented as mean ± SEM. Panels C,D,E,F,G: Unpaired t-test. Panels H&I: Repeated measures two-way ANOVA with Bonferroni post hoc test; Panels F&K: Pearson’s correlation with coefficients (r²) reported; Panels K,M,O,P: Mixed-effects analysis with Bonferroni post hoc test. Sample size: **N** = 3 - 5 mice per group (Panels C,D,E,F,G); **N** = 5 - 14 mice per group (Panels H,I,J,K,M,O,P,Q). *p < 0.05, **p < 0.01, ***p < 0.001, ****p < 0.0001; ns, not significant. Full statistical results are provided in **Extended Data Table 1**.

We next assessed signaled active avoidance in SIStag.TSC and SIStag.WT mice. Relative to controls, SIStag.TSC mice showed heightened avoidance (**Fig. 2H**) and reduced avoidance latency (**Fig. 2I**) from the first training session onward. The inverse relationship between avoidance and mean latency was robust in both genotypes (**Fig. 2J**). Because more efficient avoidance is not by itself evidence of dysfunction, we examined whether responding remained confined to the conditions that licensed it. Shuttling outside the CS epoch increased progressively across sessions in SIStag.TSC mice, whereas SIStag.WT controls maintained stable, low rates (**Fig. 2K**). Mutant shuttling is therefore not restricted to the CS but generalizes across epochs, escalating in the absence of any threat cue. CS-evoked freezing declined across days in SIStag.TSC mice relative to controls (**Fig. 2L, M**), shifting the balance from reactive toward proactive coping (**Fig. 2N**). Epoch-resolved analysis showed reduced freezing during baseline and postAR epochs on successful trials in SAA2 and SAA3 (**Fig. 2O**), and reduced freezing across baseline, CS, and postUS epochs on failed trials (**Fig. 2P**). Critically, SIStag.TSC mice lacked the positive correlation between CS-evoked freezing and avoidance latency during SAA1 that is present in controls (**Fig. 2Q**). Freezing was expressed and varied across trials but no longer predicted the latency to shuttle. The competitive phase is thus not truncated, as after social isolation alone but absent from the outset: shuttling is released from the Pavlovian response that normally gates it.

Together, these results link TSC/mTORC1 mediated autophagic suppression within stress-recruited prefrontal ensembles to a loss of response competition and the emergence of avoidance that escalates beyond the conditions that license it.

### 3. SIS tagged neurons remain cue-driven and lose action- and outcome-related signals after Tsc2 haploinsufficiency

To determine whether stress-recruited mPFC neurons carry behaviorally relevant signals during avoidance learning, we co-injected AAV5-cFos-CreERT2^44^ and AAV9-hSyn-DIO-GCaMP8m^54^ bilaterally into the mPFC, yielding robust GCaMP expression restricted to SIS-tagged neurons (**Fig. 3A**). Tsc2-floxed and wild-type mice were used to generate SIStag.TSC.GCaMP8m and SIStag.WT.GCaMP8m groups, respectively. Bulk calcium activity was recorded through a left mPFC-implanted optic ferrule by in vivo fiber photometry, using 470 nm excitation with a 405 nm isosbestic channel to correct for motion artifact. In both genotypes, tagged neurons showed reliable trial-by-trial responses to the CS, US and AR across failed and successful trials (**Fig. 3B, C**).

**Figure 3:**
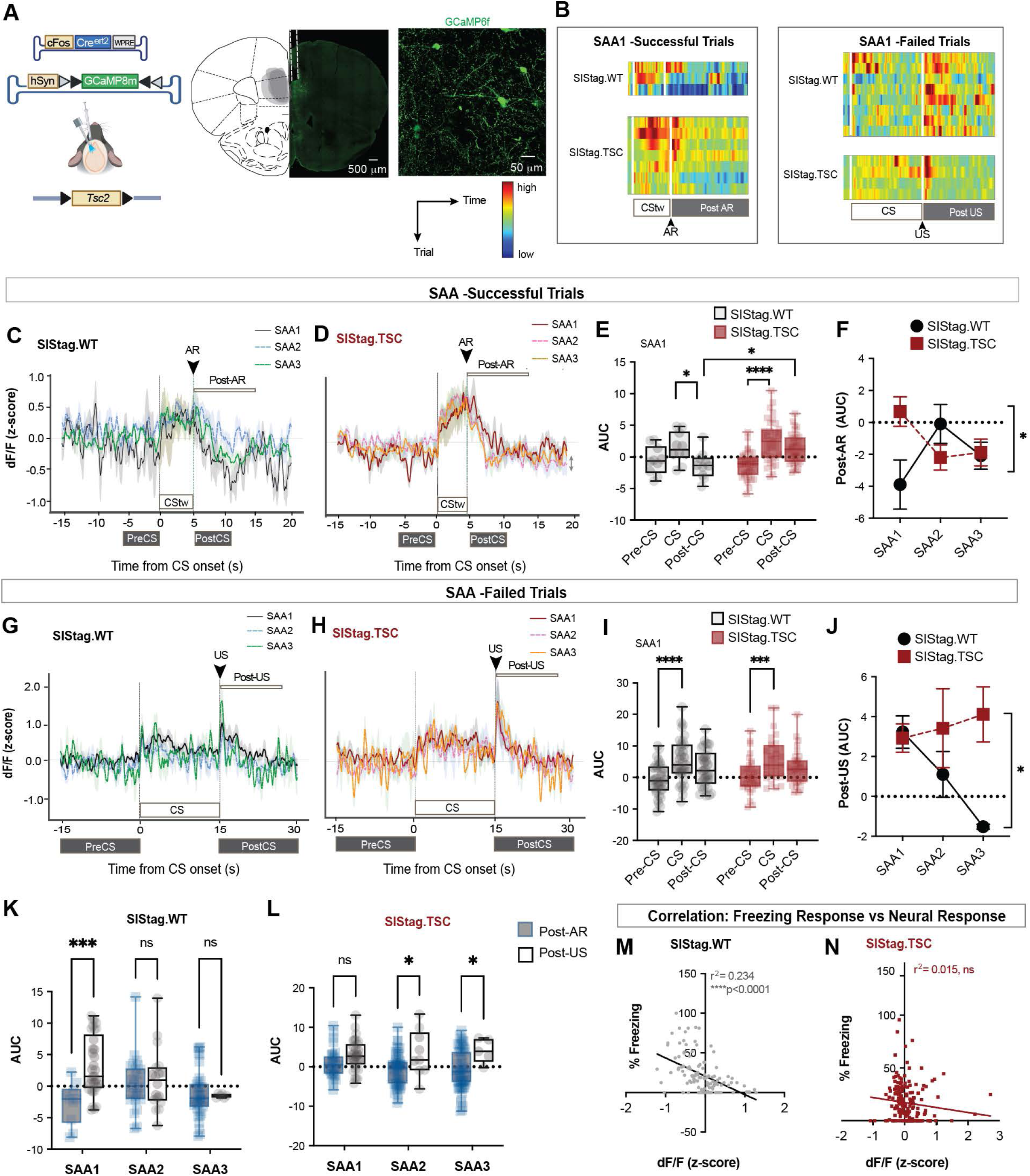
SIS tagged neurons remain cue-driven and lose action- and outcome-related signals after Tsc2 haploinsufficiency. **A.** Left, genetic strategy for activity-dependent expression of GCaMP8m in the social isolation ensemble and conditional haploinsufficiency of *Tsc2* by co-injecting AAV5-cFos-CreERT2 and AAV9-hSyn-DIO-GCaMP8m into the medial prefrontal cortex (mPFC) of *Tsc2* floxed (SIStag.TSC) and wild-type (SIStag.WT) mice. Right, representative image of GCaMP8m expression in stress-activated mPFC neurons. **B.** Heatmaps of calcium activity in stress-activated neurons during successful and failed trials of the first signaled active avoidance session (SAA1) in SIStag.WT and SIStag.TSC mice. **C.** Overlay of peri-event time histograms (PETHs) during successful trials for SIStag.WT mice across SAA1 (grey), SAA2 (dotted blue), and SAA3 (green). Time-warped conditioned stimulus (CStw), pre-CS, and post-CS epochs are indicated, with pre-CS and post-CS epochs matched to the duration of the CStw. **D.** Overlay of PETHs during successful trials for SIStag.TSC mice across SAA1 (grey), SAA2 (dotted blue), and SAA3 (green). CStw, pre-CS, and post-CS epochs are indicated. **E.** Area under the curve (AUC) of calcium responses during the pre-CS, CS, and post-CS epochs of successful SAA1 trials in SIStag.WT (black) and SIStag.TSC (red) mice. SIStag.WT mice exhibited reduced activity during the post-CS epoch relative to the CS, whereas SIStag.TSC mice displayed enhanced CS responses relative to the pre-CS epoch with no difference between the CS and post-CS epochs. **F.** Post-AR AUC across SAA1–SAA3 in SIStag.WT (black) and SIStag.TSC (red) mice. SIStag.WT mice exhibited an initially negative post-AR response that returned toward baseline during training, whereas SIStag.TSC mice showed higher initial post-AR activity that progressively decreased across sessions. **G.** Overlay of PETHs during failed trials for SIStag.WT mice across SAA1 (grey), SAA2 (dotted blue), and SAA3 (green). CStw, pre-CS, and post-CS epochs are indicated. **H.** Overlay of PETHs during failed trials for SIStag.TSC mice across SAA1 (grey), SAA2 (dotted blue), and SAA3 (green). CStw, pre-CS, and post-CS epochs are indicated. **I.** AUC of calcium responses during the pre-CS, CS, and post-CS epochs of failed SAA1 trials in SIStag.WT (black) and SIStag.TSC (red) mice. Both groups exhibited increased CS responses relative to the pre-CS epoch, with no significant differences between the CS and post-CS epochs. **J.** Post-US response AUC across SAA1–SAA3 in SIStag.WT (black) and SIStag.TSC (red) mice. SIStag.WT mice exhibited progressively reduced post-US responses across training, whereas SIStag.TSC mice maintained elevated post-US responses. **K.** Discrimination between successful and failed trials in SIStag.WT mice. Outcome-related activity distinguished successful from failed trials during SAA1 but not during later training sessions. **L.** Discrimination between successful and failed trials in SIStag.TSC mice. Outcome-related discrimination was absent during SAA1 but emerged during SAA2 and SAA3, with larger post-US responses during failed trials. **M.** Correlation between CS-evoked freezing and calcium activity (z-score) in SIStag.WT mice demonstrating a significant negative correlation. **N.** Correlation between CS-evoked freezing and calcium activity (z-score) in SIStag.TSC mice showing no significant correlation. **Statistical analyses**: Data are presented as mean ± SEM. Panels E,I,J,K,L: Mixed-effects analysis with Bonferroni post hoc test; Panel F: Repeated measures two-way ANOVA with Bonferroni post hoc test; Panels M&N: Pearson’s correlation with coefficients (r²) reported. Sample size: **N** = 4 - 6 mice per group. *p < 0.05, **p < 0.01, ***p < 0.001, ****p < 0.0001; ns, not significant. Full statistical results are provided in **Extended Data Table 1**.

To accommodate variable CS durations across successful trials, we linearly time-warped each trial to the global median CS duration ^55^. In SIStag.WT ensembles, CS-aligned activity on successful trials did not rise above pre-CS levels but was followed by pronounced suppression after the avoidance response, such that post-CS activity fell below the CS epoch (**Fig. 3D**). SIStag.TSC ensembles showed the converse: a CS-evoked increase over pre-CS with no subsequent suppression, remaining elevated through the post-CS epoch and differing from controls (**Fig. 3E**). Epoch-resolved analysis revealed that the two genotypes reached these profiles along opposite trajectories (**Extended Data Fig. 1A**). In SIStag.WT mice, CS-epoch activity exceeded post-CS but not pre-CS during SAA1, was indistinguishable from both epochs during SAA2, and rose above both only by SAA3. SIStag.TSC ensembles, by contrast, showed CS activity above pre-CS from SAA1 onward and maintained this elevation through SAA2 and SAA3 (**Fig. 3F; Extended Data Fig. 1B, C**). Control ensembles thus acquire a CS response over training, whereas mutant ensembles are cue-driven from the first session. The post-AR signature followed a corresponding inversion. In SIStag.WT mice it was strongly negative during SAA1 and rose toward zero with training, whereas in SIStag.TSC mice it began near zero and became negative only by SAA2-SAA3 (**Fig. 3G**); the genotypes diverged most sharply during SAA1, when control suppression was deepest and mutant suppression absent. The avoidance signature is therefore not simply lost in mutants but temporally displaced - present in controls while responding is still being acquired, and emerging in mutants only once responding is established.

On failed trials, both genotypes showed robust CS and US-evoked responses (**Fig. 3H, I; Extended Data Fig. 1D**). CS activity was significantly elevated over pre-CS in both genotypes at SAA1 (**Fig. 3J**) and SAA2 (**Extended Data Fig. 1E**), but not at SAA3 (**Extended Data Fig. 1F**). The genotypes diverged most clearly in the response to the shock itself: the post-US response in SIStag.WT mice declined progressively across sessions and fell below baseline by SAA3, whereas in SIStag.TSC mice it remained elevated and did not habituate (**Fig. 3K**).

Comparing post-AR and post-US responses directly, SIStag.WT ensembles distinguished successful from failed trials during SAA1 but not thereafter (**Fig. 3L**). SIStag.TSC ensembles showed the opposite profile, failing to discriminate during SAA1 and separating the two trial types only in SAA2 and SAA3 (**Fig. 3M**). Outcome-related information is thus present in control ensembles precisely during the phase when freezing and shuttling compete and appears in mutants only once responding has become stereotyped. If the ensemble arbitrates between freezing and shuttling, its activity should track the competing response. We found that in SIStag.WT mice, post-CS calcium activity was strongly negatively correlated with freezing (r^2^ = 0.234, p < 0.0001), consistent with reciprocal control of the two defensive responses (**Fig. 3N**). This relationship was absent in SIStag.TSC mice (r^2^ = 0.015, ns; **Fig. 3O**). The mutant ensemble therefore no longer tracks the Pavlovian response it must otherwise override - a neural counterpart to the behavioral decoupling of freezing from avoidance latency.

Together, these results indicate that Tsc2 haploinsufficiency in stress-recruited ensembles does not silence prefrontal activity but reorganizes what it represents: the CS response is amplified, the post-action signature and outcome discrimination are shifted out of the acquisition phase, and the reciprocal relationship with freezing is lost entirely.

### 4. Social isolation accelerates avoidance memory whereas Tsc2 haploinsufficiency unbinds it from the cue

We assessed the associative memory linking the CS to the US by testing retrieval under non-reinforced conditions in a novel context at two timepoints: 24 h after SAA1 (LTM1) and 24 h after SAA3 (LTM2) (**Fig. 4A**). Mice received five CS presentations, with no shock delivered on failed trials, allowing memory expression to be measured after one versus three days of training. We first compared group-housed and socially isolated wild-type mice. At LTM1, socially isolated (SIS) mice showed markedly higher CS-evoked avoidance than group-housed (GH) controls; by LTM2, the two groups were indistinguishable (**Fig. 4B**). Avoidance latency followed the same pattern, being shorter in SIS mice at LTM1 and comparable between groups at LTM2, once additional training had brought group-housed animals to the same level (**Fig. 4C**). Shuttling outside the CS epoch was minimal and did not differ between groups (**Fig. 4D**), and CS-evoked freezing was unchanged both between groups and across timepoints (**Fig. 4E**). Epoch-resolved analysis confirmed that freezing remained low across peri-CS and CS epochs on both successful and failed trials (**Extended Data Fig. 2A, B**), and AR latency was not correlated with CS-evoked freezing in either group (**Extended Data Fig. 2C**). Social isolation therefore accelerates the consolidation of avoidance memory, producing after a single session the level of retrieval that controls reach only after three - without generalized shuttling outside the conditions that license it.

**Figure 4:**
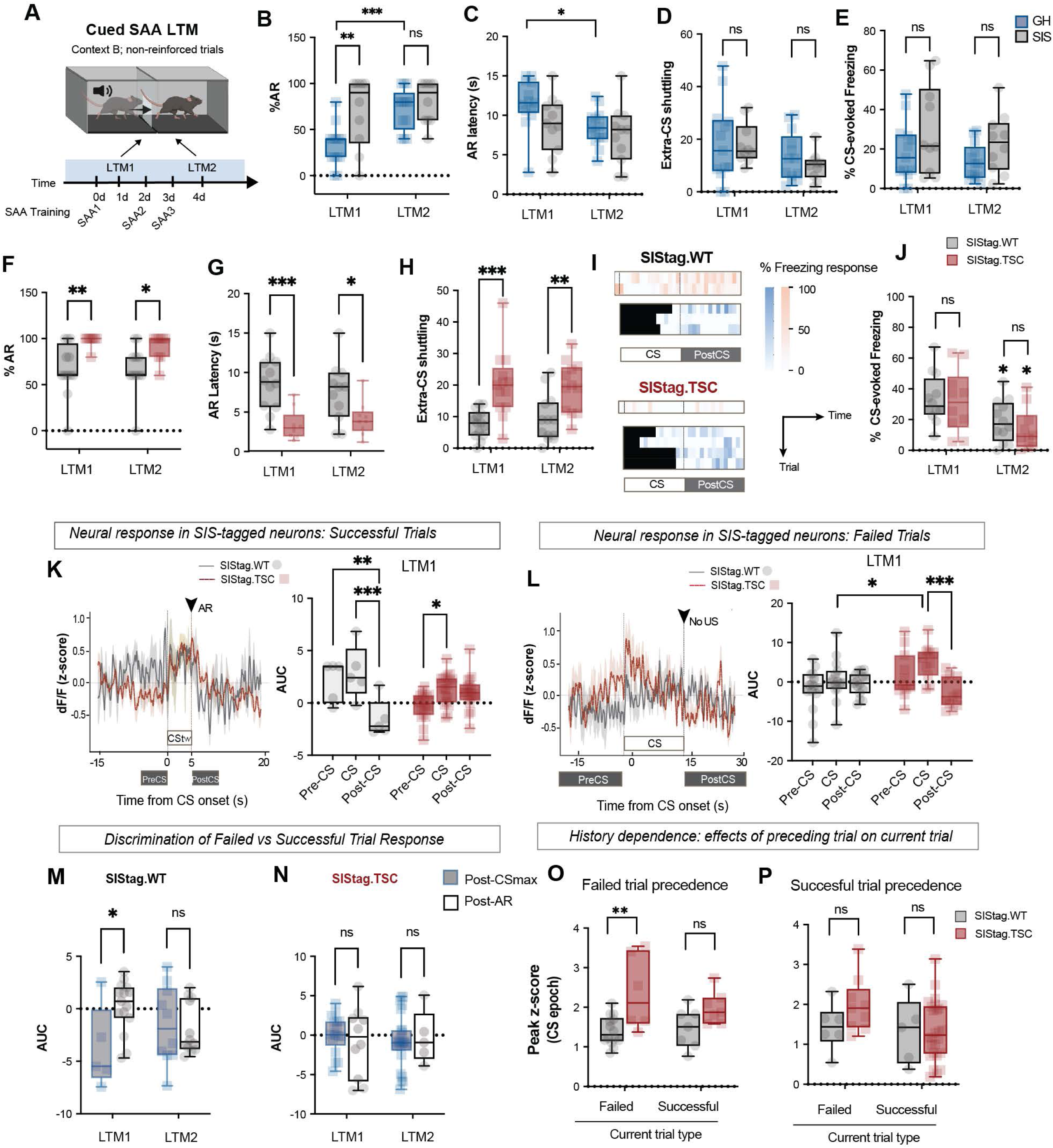
Social isolation accelerates avoidance memory whereas Tsc2 haploinsufficiency unbinds it from the cue. **A.** LTM testing paradigm and timeline. Mice underwent SAA memory retrieval in a context distinct from the training context. LTM1 was conducted 24 h after SAA1, and LTM2 was conducted 24 h after SAA3. **B.** Percentage of CS-evoked avoidance responses during LTM1 and LTM2. SIS (grey) mice exhibited increased avoidance responses relative to GH (blue) mice during LTM1, with no group differences during LTM2. **C.** Latency to initiate the avoidance response during LTM1 and LTM2. SIS mice exhibited equivalent avoidance latencies as GH mice during LTM1 and LTM2. **D.** Extra-CS shuttling during LTM1 and LTM2. No significant differences were observed between GH and SIS mice. **E.** Percentage of CS-evoked freezing during LTM1 and LTM2. No significant differences were observed between GH and SIS mice. **F.** Percentage of CS-evoked avoidance responses during LTM1 and LTM2. SIStag.TSC mice exhibited increased avoidance responses relative to SIStag.WT controls during both memory tests. **G.** Latency to initiate the avoidance response during LTM1 and LTM2. SIStag.TSC mice exhibited shorter avoidance latencies than SIStag.WT mice during both memory tests. **H.** Extra-CS shuttling during LTM1 and LTM2. SIStag.TSC mice exhibited increased extra-CS shuttling relative to SIStag.WT controls during both memory tests. **I.** Representative freezing heatmaps during failed (top) and successful (bottom) trials aligned to the CS and post-US or post-AR periods. **J.** Percentage of CS-evoked freezing during LTM1 and LTM2. No significant differences were observed between SIStag.WT and SIStag.TSC mice. **K.** Left, overlay of PETHs during successful trials for LTM1. CStw, pre-CS, and post-CS epochs are indicated. Right, AUC of calcium responses during the pre-CS, CS, and post-CS epochs of successful LTM1 trials. Both SIStag.WT and SIStag.TSC mice exhibited increased activity during the CS epoch relative to the pre-CS epoch. SIStag.WT mice exhibited reduced post-CS responses relative to the CS epoch, whereas SIStag.TSC mice exhibited no significant differences between the CS and post-CS epochs. **L.** Left, overlay of PETHs during failed trials for LTM1. CStw, pre-CS, and post-CS epochs are indicated. Right, AUC of calcium responses during the pre-CS, CS, and post-CS epochs of failed LTM1 trials. SIStag.WT mice exhibited no significant differences across epochs, whereas SIStag.TSC mice exhibited reduced post-CS activity relative to the CS epoch and greater CS responses than SIStag.WT mice. **M.** Discrimination between successful and failed trials in SIStag.WT mice. Outcome-related activity distinguished successful from failed trials during LTM1 but not LTM2. **N.** Discrimination between successful and failed trials in SIStag.TSC mice. Outcome-related discrimination was absent during both LTM sessions. **O.** Peak calcium responses during trials preceded by a failed trial. SIStag.TSC mice exhibited greater peak responses during failed trials than SIStag.WT mice, with no significant group differences during successful trials. **P.** Peak calcium responses during trials preceded by a successful trial. No significant differences were observed between SIStag.WT and SIStag.TSC mice during either successful or failed trials. **Statistical analyses**: Data are presented as mean ± SEM. Panels B,C,D,E,F,H,J,K,L,M,N: Mixed-effects analysis with Bonferroni post hoc test; Panel G,O,P: Two-way ANOVA with Bonferroni post hoc test. Sample size: **N** = 9 - 14 mice per group (Panels B-J); **N** = 4 - 6 mice per group (Panels K-P). *p < 0.05, **p < 0.01, ***p < 0.001, ****p < 0.0001; ns, not significant. Full statistical results are provided in **Extended Data Table 1**.

We next asked whether Tsc2 haploinsufficiency in stress-recruited ensembles produced a comparable acceleration, or a qualitatively different retrieval profile. SIStag.TSC mice showed enhanced CS-evoked avoidance relative to SIStag.WT controls at both LTM1 and LTM2 (**Fig. 4F**), with correspondingly shorter avoidance latencies at both timepoints (**Fig. 4G**). Unlike the isolation-alone comparison, the two genotypes did not converge with further training. Shuttling outside the CS epoch was also elevated in SIStag.TSC mice at both LTM1 and LTM2 (**Fig. 4H**), demonstrating that mutant responding is not confined to the cue even in a novel context and in the absence of reinforcement. CS-evoked freezing was indistinguishable between genotypes (**Fig. 4I, J**), as was epoch-resolved freezing on both successful and failed trials (**Extended Data Fig. 3A, B**), and AR latency was uncorrelated with CS-evoked freezing in either group (**Extended Data Fig. 3C**). Tsc2 haploinsufficiency in stress ensembles therefore does not simply accelerate avoidance memory as isolation does but yields retrieval in which shuttling has generalized beyond the CS that licenses it - a dissociation that persists rather than resolving with training.

To determine how stress ensembles represent retrieval, we recorded calcium activity by fiber photometry during LTM1 and LTM2. On successful trials at LTM1, SIStag.WT ensembles showed a modest CS-evoked increase followed by suppression of activity in the post-CS epoch immediately after the avoidance response. SIStag.TSC ensembles showed the converse: a significant CS-evoked increase over pre-CS, which controls lacked, and no subsequent post-AR suppression (**Fig. 4K; Extended Data Fig. 5A, B**). By LTM2, control ensembles showed no significant modulation across any epoch, whereas mutant ensembles retained a CS response (**Extended Data Fig. 5C**). The post-AR response was thus strongly divergent at LTM1 - deeply negative in controls, near zero in mutants - and converged by LTM2 (**Extended Data Fig. 5D**).

On failed trials, SIStag.TSC ensembles showed a heightened CS response that was absent in controls, followed by suppression of activity when the expected US was omitted (**Fig. 4L; Extended Data Fig. 5E, F**). So, the mutant ensemble responds to a violated expectation but not to the outcome distinction. SIStag.WT ensembles showed negligible modulation during either the CS or post-CS_max_ epochs at both timepoints (**Extended Data Fig. 5G**), and the post-CS_max_ response did not differ between genotypes (**Extended Data Fig. 5H**).

Comparing post-AR and post-CS_max_ responses directly, SIStag.WT ensembles discriminated successful from failed trials at LTM1 but not at LTM2 (**Fig. 4M**), whereas SIStag.TSC ensembles failed to discriminate at either timepoint (**Fig. 4N**). Outcome-related information is therefore present in control ensembles during early retrieval and disappears once performance is asymptotic; but is never detectable in mutants. Finally, we asked whether ensemble activity was influenced by trial history. In SIStag.TSC mice, the peak CS response on a failed trial was elevated when the preceding trial had also been a failure, an effect absent in controls; no such dependence was observed when the current trial was successful (**Fig. 4O**). Preceding successful trials had no effect on either trial type in either genotype (**Fig. 4P**). Mutant ensembles are thus selectively sensitive to recent failure, amplifying the cue response while remaining unable to distinguish outcomes at the time they occur.

Together, these results distinguish two forms of altered avoidance memory: social isolation accelerates consolidation without unbinding the response from its cue, whereas Tsc2 haploinsufficiency in stress ensembles produces retrieval in which shuttling generalizes beyond the CS and the ensemble amplifies the cue-evoked neural response - including after recent failure - while never signaling which outcome the animal obtained.

### 5. mTORC1 inhibition in stress ensembles restores maladaptive avoidance but not Pavlovian freezing

Because Tsc2 haploinsufficiency in stress ensembles suppressed autophagy downstream of hyperactive mTORC1, we asked whether restoring normative mTORC1 signaling would rescue the behavioral phenotype, using two independent strategies: cell-type-specific knockdown of Raptor within SIStag neurons, and pharmacological inhibition of mTORC1 with rapamycin (**Fig. 5A**). Raptor is an obligate structural component of mTORC1, so shRNA-mediated knockdown depletes the complex in a genetically defined population. We co-injected AAV5-cFos-CreERT2, AAV-CAG-DIO-tdTomato, and a pSICO-based U6-shRNA[Raptor] lentivirus bilaterally into the mPFC of Tsc2-floxed mice, restricting knockdown to SIS-tagged cells (**Fig. 5B**). Immunohistochemistry confirmed reduced Raptor immunoreactivity within tdTomato+ ensemble neurons relative to SIStag.TSC mice lacking the shRNA (**Fig. 5C**). Mice were singly housed 21 days after surgery, injected with 4-OHT on the first day of isolation, and trained beginning 10 days later (**Fig. 5D**). During reinforced training, SIStag.TSC+shRaptor mice acquired avoidance at rates and latencies indistinguishable from SIStag.WT controls (**Fig. 5E, F**). Raptor knockdown in TSC mutants did not, however, rescue selective response to the cue: shuttling outside the CS epoch increased progressively across sessions and exceeded control levels (**Fig. 5G**), and CS-evoked freezing remained reduced throughout training (**Fig. 5H**). At LTM retrieval, Raptor knockdown reversed the phenotype. SIStag.TSC mice showed elevated CS-evoked avoidance and shortened latency at LTM1; both were restored to levels indistinguishable from SIStag.WT by Raptor knockdown (**Fig. 5I, J**), with all groups converging by LTM2. Extra-CS shuttling, elevated in SIStag.TSC mice at both timepoints, was likewise no longer distinguishable from controls following knockdown (**Fig. 5K**). CS-evoked freezing did not differ among groups (**Fig. 5L**).

**Figure 5:**
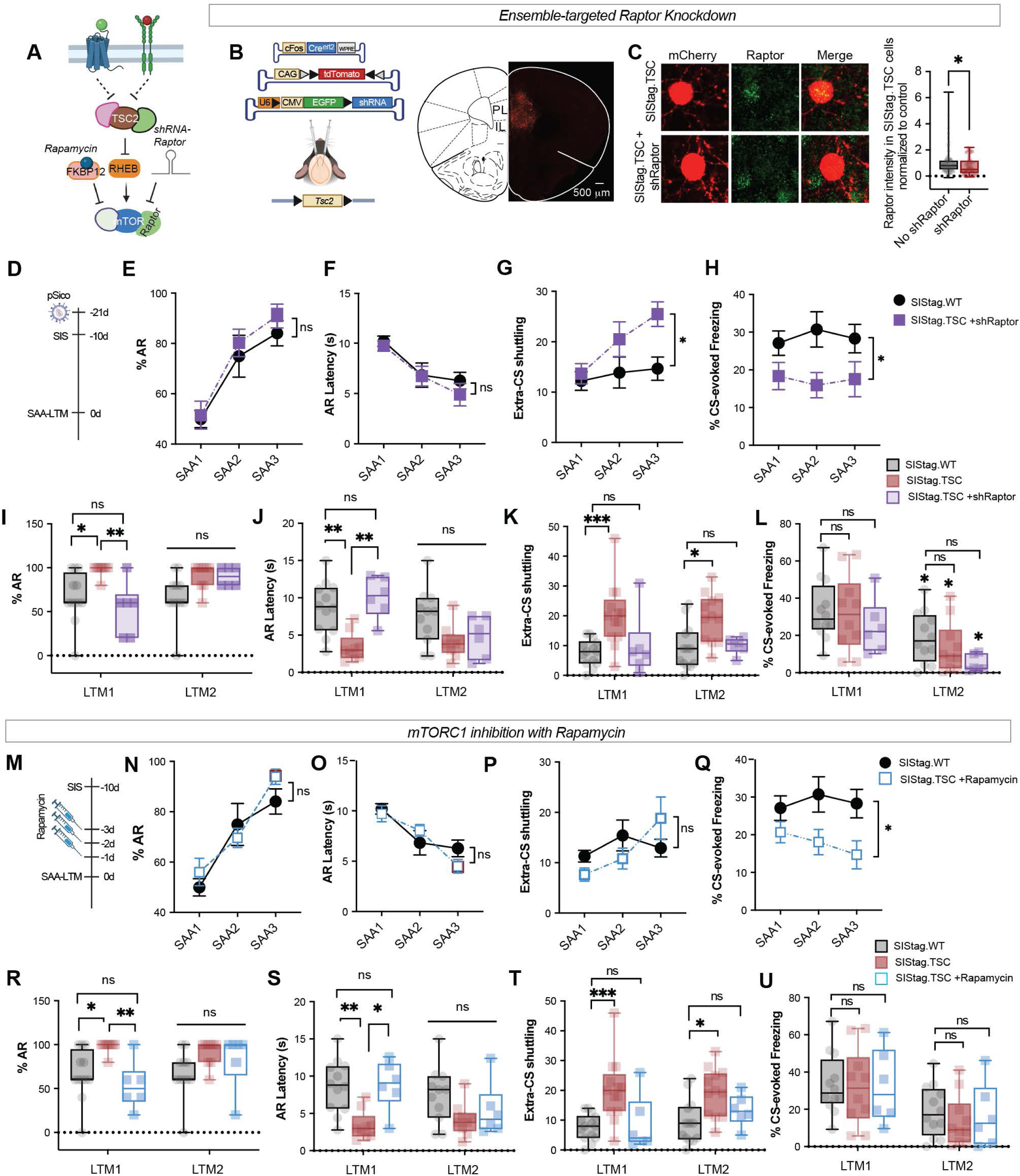
mTORC1 inhibition in stress ensembles restores maladaptive avoidance but not Pavlovian freezing. **A.** Schematic illustrating the rescue strategies using Raptor knockdown and rapamycin treatment. **B.** Left, genetic strategy for activity-dependent Raptor knockdown by co-injecting AAV5-cFos-CreERT2, AAV-CAG-DIO-tdTomato, and a pSICO-based U6-shRNA[Raptor] lentivirus bilaterally into the mPFC of *Tsc2*-floxed mice. Right, representative image of tdTomato-labeled social isolation ensemble neurons expressing shRaptor in the mPFC. Scale bar = 500 μm. **C.** Left, representative images of Raptor immunofluorescence in SIStag.WT and SIStag.TSC+shRaptor neurons. Scale bar = 50 μm. Right, quantification of Raptor fluorescence intensity demonstrating a significant reduction in SIStag.TSC+shRaptor (purple) relative to SIStag.WT (black). **D.** Timeline for viral surgery, social isolation, and behavioral testing. **E.** Percentage of CS-evoked avoidance responses across SAA1–SAA3. No significant differences were observed between SIStag.TSC+shRaptor (purple) and SIStag.WT (black) mice. **F.** Latency to initiate the avoidance response during SAA1–SAA3. No significant differences were observed between SIStag.TSC+shRaptor and SIStag.WT mice. **G.** Extra-CS shuttling during SAA training. SIStag.TSC+shRaptor mice exhibited increased extra-CS shuttling relative to SIStag.WT controls. **H.** Percentage of CS-evoked freezing across SAA1–SAA3. SIStag.TSC+shRaptor mice exhibited reduced freezing relative to SIStag.WT controls, indicating that freezing behavior was not rescued. **I.** Percentage of CS-evoked avoidance responses during LTM1 and LTM2. SIStag.TSC+shRaptor mice exhibited avoidance responses comparable to SIStag.WT controls during both memory tests. **J.** Latency to initiate the avoidance response during LTM1 and LTM2. SIStag.TSC+shRaptor mice exhibited avoidance latencies comparable to SIStag.WT controls during both memory tests. **K.** Extra-CS shuttling during LTM1 and LTM2. SIStag.TSC+shRaptor mice exhibited extra-CS shuttling comparable to SIStag.WT controls during both memory tests. **L.** Percentage of CS-evoked freezing during LTM1 and LTM2. No significant differences were observed between groups during either memory test. **M.** Timeline for rapamycin treatment. Mice received rapamycin for 3 days following social isolation before SAA training. **N.** Percentage of CS-evoked avoidance responses across SAA1–SAA3. No significant differences were observed between SIStag.TSC+Rapamycin (blue) and SIStag.WT (black) mice. **O.** Latency to initiate the avoidance response during SAA1–SAA3. No significant differences were observed between SIStag.TSC+Rapamycin and SIStag.WT mice. **P.** Extra-CS shuttling during SAA1–SAA3. SIStag.TSC+Rapamycin mice exhibited extra-CS shuttling comparable to SIStag.WT controls. **Q.** Percentage of CS-evoked freezing across SAA1–SAA3. SIStag.TSC+Rapamycin mice exhibited reduced freezing relative to SIStag.WT controls, indicating that freezing behavior was not rescued. **R.** Percentage of CS-evoked avoidance responses during LTM1 and LTM2. SIStag.TSC+Rapamycin mice exhibited avoidance responses comparable to SIStag.WT controls during both memory tests. **S.** Latency to initiate the avoidance response during LTM1 and LTM2. SIStag.TSC+Rapamycin mice exhibited avoidance latencies comparable to SIStag.WT controls during both memory tests. **T.** Extra-CS shuttling during LTM1 and LTM2. SIStag.TSC+Rapamycin mice exhibited extra-CS shuttling comparable to SIStag.WT controls during both memory tests. **U.** Percentage of CS-evoked freezing during LTM1 and LTM2. No significant differences were observed between groups during either memory test. **Statistical analyses**: Data are presented as mean ± SEM. Panel C: Unpaired t-test. Panels E-U: Mixed-effects analysis with Bonferroni post hoc test. Sample size: **N** = 5 - 6 mice per group (Panel C); **N** = 6 - 14 mice per group (Panels E-U). *p < 0.05, **p < 0.01, ***p < 0.001, ****p < 0.0001; ns, not significant. Full statistical results are provided in **Extended Data Table 1**.

We next asked whether pharmacologically reinstating restraint on mTORC1 would reproduce this profile, administering rapamycin intraperitoneally on the three days preceding training (**Fig. 5M**). As with Raptor knockdown, avoidance responses and latencies during training were comparable to controls (**Fig. 5N, O**). Unlike shRaptor, rapamycin also normalized extra-CS shuttling during training (**Fig. 5P**), while CS-evoked freezing remained reduced relative to controls (**Fig. 5Q**). At memory retrieval, rapamycin restored CS-evoked avoidance responses and avoidance latency to control levels at LTM1 (**Fig. 5R, S**), with convergence by LTM2. Extra-CS shuttling, elevated in SIStag.TSC mice at both timepoints, was returned to control levels by rapamycin (**Fig. 5T**), and CS-evoked freezing was unaffected (**Fig. 5U**).

Two independent manipulations of mTORC1 signaling therefore converge on the same pattern: both restore the vigor of avoidance and neither restores the Pavlovian freezing response, while rapamycin additionally confines responding to the cue. The maladaptive features of avoidance in SIStag.TSC mice - its excessive vigor and its escape from the conditions that license it - are thus mTORC1-dependent and reversible, whereas the loss of CS-evoked freezing is not. Suppression of the Pavlovian response and dysregulation of the instrumental one are therefore separable consequences of Tsc2 haploinsufficiency.

### 6. The avoidance phenotype is specific to threat-based decision making and requires social isolation

To determine whether Tsc2 haploinsufficiency in mPFC stress ensembles affects other prefrontal cortex dependent behaviors, we tested a battery of assays: open field and elevated plus maze for innate anxiety, self-grooming for stereotypy, and novel object recognition for long-term memory that is neither associative nor threat-related. SIStag.TSC mice habituated to the open field and explored comparably to SIStag.WT controls, with equivalent locomotion across the session and equivalent total distance travelled (**Extended Data Fig. 6A-C**). Thigmotaxis, measured as the ratio of center to total distance, did not differ between genotypes (**Extended Data Fig. 6D**). In the elevated plus maze, the two groups showed comparable open-arm duration, open-arm entries, and latency to enter the open arm (**Extended Data Fig. 6E-H**). Tsc2 haploinsufficiency in stress ensembles, combined with the experience of social isolation, therefore does not precipitate innate anxiety. Grooming duration and latency to groom were likewise unaffected (**Extended Data Fig. 6I-K**), indicating no innate compulsive or stereotyped responding.

We next tested long-term memory for a neutral stimulus. Mice were habituated to two identical objects across three days, and one was replaced with a novel object at test (**Fig. 6A**). Interaction was quantified from DeepLabCut-based pose estimation, scoring frame-by-frame occupancy of the head within a 1 inch (∼2.54 cm) zone surrounding each object (**Fig. 6B**), and recognition memory was expressed as a novelty preference index, PI = (NO − FO) / (NO + FO). Both genotypes preferred the novel object, with equivalent preference indices (**Fig. 6C**) and equivalent latencies to first interact with it (**Fig. 6D**). Tsc2 haploinsufficiency in stress ensembles therefore leaves retrieval of a non-associative, neutral memory intact.

**Figure 6:**
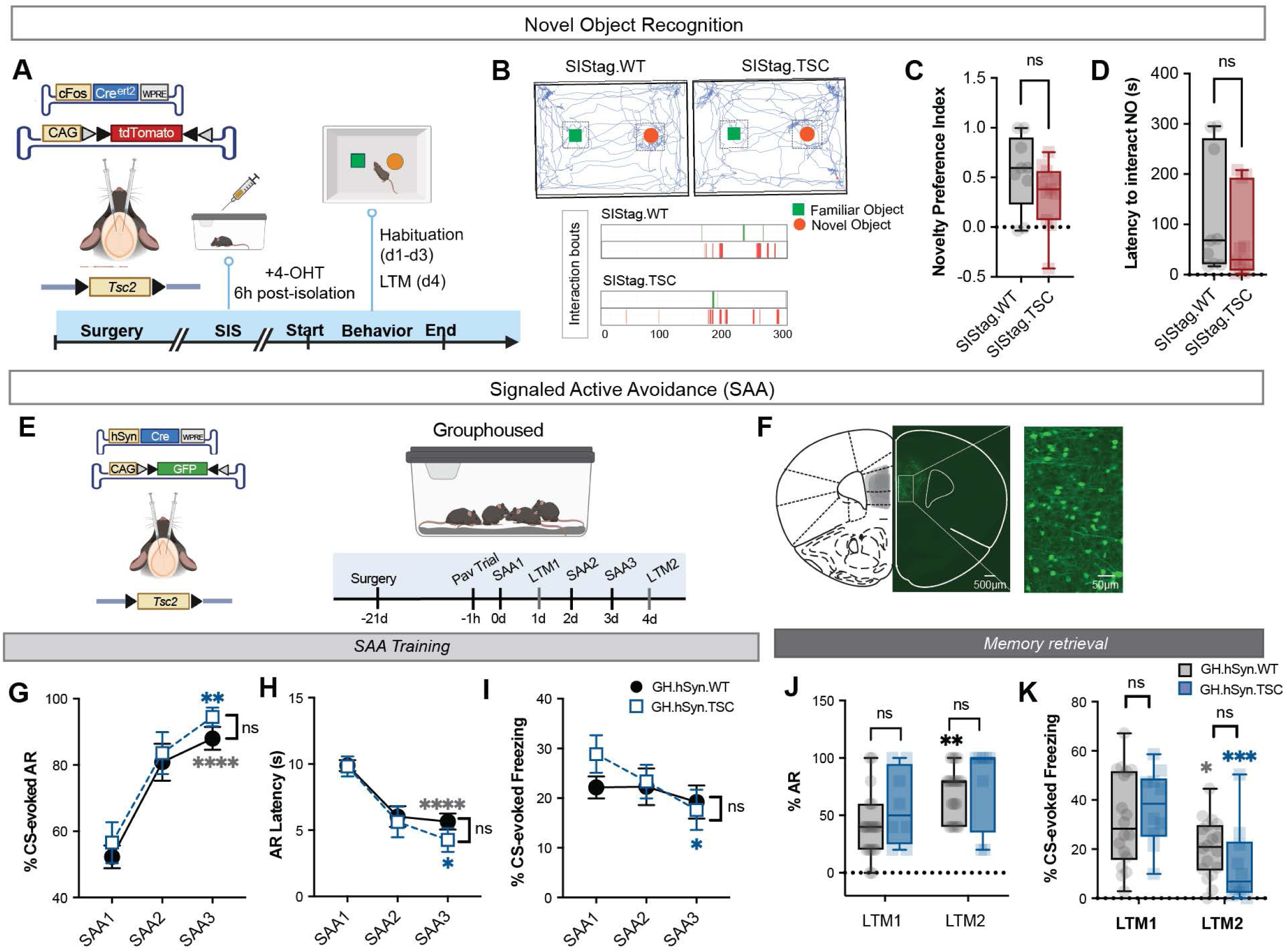
The avoidance phenotype is specific to threat-based decision making and requires social isolation. **A.** Timeline for the novel object recognition (NOR) task following *Tsc2* haploinsufficiency in stress-activated neurons. Mice were habituated to the familiar objects for 3 days before testing 24 h later. **B.** Upper, representative heatmaps showing object exploration by SIStag.WT and SIStag.TSC mice during the NOR test. Lower, object interaction bouts over time for familiar objects (FO) and the novel object (NO). **C.** Novelty preference index. No significant differences were observed between SIStag.WT and SIStag.TSC mice. **D.** Latency to interact with the NO. No significant differences were observed between SIStag.WT and SIStag.TSC mice. **E.** Left, genetic strategy for random haploinsufficiency of *Tsc2* in mPFC neurons by co-injecting AAV9-hSyn-Cre and AAV9-CAG-DIO-GFP bilaterally into the mPFC of *Tsc2*-floxed or wild-type mice. Right, experimental timeline showing SAA training following viral surgery in GH mice. **F.** Representative images of GFP-labeled mPFC neurons following *Tsc2* haploinsufficiency. Scale bar = 500 μm, 50 μm. **G.** Percentage of CS-evoked avoidance responses across SAA1–SAA3. No significant differences were observed between GH.hSyn.WT (blue) and GH.hSyn.TSC (black) mice. **H.** Latency to initiate the avoidance response during SAA1–SAA3. No significant differences were observed between GH.hSyn.WT and GH.hSyn.TSC mice. **I.** Percentage of CS-evoked freezing across SAA1–SAA3. No significant differences were observed between GH.hSyn.WT and GH.hSyn.TSC mice. **J.** Percentage of CS-evoked avoidance responses during LTM1 and LTM2. No significant differences were observed between GH.hSyn.WT and GH.hSyn.TSC mice during either memory test. **K.** Percentage of CS-evoked freezing during LTM1 and LTM2. No significant differences were observed between GH.hSyn.WT and GH.hSyn.TSC mice during either memory test. **Statistical analyses**: Data are presented as mean ± SEM. Panel C&D: Unpaired t-test. Panels G-K: Mixed-effects analysis with Bonferroni post hoc test. Sample size: **N** = 8 - 20 mice per group. *p < 0.05, **p < 0.01, ***p < 0.001, ****p < 0.0001; ns, not significant. Full statistical results are provided in **Extended Data Table 1**.

The avoidance phenotype thus appears specific to threat-based decision making. We next asked whether it also requires the stress experience itself, or whether Tsc2 loss in prefrontal neurons is sufficient to produce it in the absence of adversity. We co-injected AAV9-hSyn-Cre and AAV9-CAG-DIO-GFP bilaterally into the mPFC of wild-type or Tsc2-floxed mice, deleting Tsc2 across approximately 85 neurons - comparable to the ∼60 SIStag neurons targeted in the same regions earlier - distributed through PL and IL of each hemisphere, and trained animals on SAA under continuous group housing (**Fig. 6E, F**). Both genotypes acquired avoidance normally, with comparable avoidance rates and latencies across all three training sessions (**Fig. 6G,H**) and comparable CS-evoked freezing (**Fig. 6I**). Retrieval was likewise unaffected, with no genotype difference in CS-evoked avoidance or freezing at either LTM1 or LTM2 (**Fig. 6J,K**). Extra-CS shuttling was unchanged during reinforced trials of SAA training and non-reinforced trials of LTM1 and LTM2 (**Extended Data Fig. 7A, B**).

Together, these findings indicate that Tsc2 haploinsufficiency in stress-recruited mPFC neurons selectively alters threat-based decision making, leaving innate anxiety, stereotypy, and neutral recognition memory intact. Critically, prefrontal Tsc2 haploinsufficiency in the absence of social isolation produces no avoidance phenotype at all: genetic vulnerability alone is insufficient, and the maladaptive strategy emerges only when Tsc2 haploinsufficiency coincides with the stress experience that recruits the ensemble in which it acts.

## Discussion

We asked how genetic vulnerability and environmental adversity converge to transform adaptive threat learning into pathological avoidance. Using activity-dependent targeting to access the mPFC neurons recruited during the early phase of social isolation ^19,22,43^, we find that deleting Tsc2 within this population produces avoidance that is efficient by every conventional performance measure yet escapes the conditions that license it. These animals never pass through the phase in which freezing and shuttling compete; their responding invades the extra-CS epochs and generalizes to a novel context in which no shock has ever been delivered; and the ensemble that drives them remains locked to the cue while failing to signal what the action achieved. This phenotype emerges only when the genetic lesion and the stress are combined, neither being sufficient alone.

Signaled active avoidance is unusual among threat paradigms in that success requires overcoming a defensive reflex rather than expressing one ^46,47^. The positive correlation between CS-evoked freezing and shuttle latency in control animals is the behavioral signature of that conflict: freezing imposes a cost on the latency to act ^45,56^, and the period over which the correlation persists indexes the window in which the two systems are arbitrating. Social isolation compresses that window to a single session, and Tsc2 haploinsufficiency removes it altogether. Critically, mutant animals still freeze, and their freezing still varies across trials - it simply no longer predicts when they will shuttle, so the Pavlovian response is expressed but disconnected from the instrumental one. This dissociation, rather than any change in avoidance rate, is what distinguishes the mutant strategy from a normal, efficiently acquired habit ^57,58^: control animals also stop freezing before shuttling by the end of training, but they earn that endpoint by resolving the conflict, whereas mutants arrive there without ever engaging it. More avoidance is not, by itself, evidence of dysfunction; within the task, mutants are better adapted than controls, receiving fewer shocks at no measurable cost. What renders their avoidance maladaptive is instead its independence from stimulus control: shuttling escalates progressively in the absence of any threat cue and persists in a novel context under non-reinforced conditions. This mirrors the structure of pathological avoidance in humans, which is not typically ineffective at preventing the feared outcome but rather insensitive to whether the outcome remains a threat at all ^59–61^. A cached policy is optimal in a stationary world; its pathology becomes visible only when the world changes or when the response spreads beyond the circumstances that justified it ^62,63^, which is precisely what our data capture.

Fiber photometry from the same ensembles shows that Tsc2 haploinsufficiency does not diminish prefrontal activity - it redistributes what that activity represents. Control ensembles are cue-responsive early and disengage as avoidance becomes efficient; they carry a deep post-action suppression and discriminate successful from failed trials precisely during the session in which freezing and shuttling compete, and their post-CS activity tracks freezing reciprocally. Mutant ensembles show the mirror image: a persistent CS response present from the first session, no early post-action signature, outcome discrimination that appears only once responding is stereotyped and is undetectable at retrieval, and no relationship to freezing at any point, such that the ensemble becomes louder and less informative. Two features are worth emphasizing. First, mutant activity is history-dependent in a selective way - a failed trial amplifies the cue response on the next failed trial, while successful trials carry no such influence - indicating that the ensemble registers aversive events without representing the distinction between outcomes when they occur. Second, mutant ensembles suppress activity when an expected shock is omitted, so the failure is not a general insensitivity to consequence but a specific absence of the outcome variable at the time of choice. The most parsimonious reading is that the ensemble has become a cue detector rather than an arbitrator, and that the safety signal which normally terminates a threat state - and thereby defines its boundaries ^64–66^ - is no longer generated. Without that boundary, responding is unconstrained and extends into the intervals and contexts where it does not belong.

Heterozygous Tsc2 loss in stress ensembles suppressed autophagic flux without detectably altering bulk translation, dissociating the two arms that mTORC1 normally gates in opposition ^28–30^. This dissociation was unexpected, since the canonical expectation is that mTORC1 hyperactivation elevates cap-dependent synthesis ^67,68^; it may be that heterozygous loss produces a level of deregulation to which the degradative arm is more sensitive than the synthetic one, or that changes are confined to specific transcripts that bulk AHA incorporation cannot resolve ^68^. Either way, the phenotype in these neurons is a loss of turnover rather than an excess of production - a distinction with consequences in post-mitotic neurons, where autophagy is required for receptor turnover, spine elimination, and the removal of synaptic proteins that bidirectional plasticity depends on ^31–35^. A neuron capable of synthesis but deficient in degradation constitutes a plausible substrate for a controller that acquires a policy early and cannot revise it, and the somatodendritic hypertrophy we observe is consistent with that imbalance and with the spine and dendritic phenotypes reported in other TSC models ^36–38^.

Previous work has shown that diverse chronic stressors - social defeat, restraint, foot shock – suppress autophagic flux selectively in the lateral habenula while sparing the mPFC ^69^. Chronic stress has been shown to affect mPFC through a different axis: sustained catecholamine and glucocorticoid signaling that lead to withdrawal of the BDNF-mTORC1 trophic support, remodel the actin cytoskeleton, and produce spine loss and apical dendritic retraction ^70,71^. This route lowers mTORC1 activity and yields neuronal atrophy - the mirror image of the mTORC1 hyperactivation and somatodendritic hypertrophy that heterozygous Tsc2 loss produces in these same ensembles. The two arms thus impose opposing effects on mTORC1 activity, yet these do not offset at the level that matters functionally: because chronic stress spares mPFC autophagy, the genetic lesion imposes an autophagic suppression that stress alone does not, and we propose that this uncorrected deficit in proteostatic turnover within stress-recruited ensembles drives maladaptive avoidance.

Two independent manipulations of mTORC1 - ensemble-restricted Raptor knockdown and systemic rapamycin - converge on the same outcome: both normalize avoidance vigor and restore stimulus control at retrieval, and neither restores CS-evoked freezing. The convergence of a cell-type-specific genetic manipulation and a systemic pharmacological one on the same profile argues that the rescued components are mTORC1-dependent within the ensemble itself, consistent with rapamycin rescue of learning deficits in Tsc2 haploinsufficient mice ^72,73^, while the freezing deficit, by contrast, resists both, indicating that suppression of the Pavlovian response and dysregulation of the instrumental one are separable consequences of Tsc2 loss. It is possible that mTORC1 inhibition may lower protein synthesis below the threshold required to consolidate the CS-US association ^74–76^, in which case the persistent freezing deficit is a cost of the intervention rather than an unrescued phenotype.

Deleting Tsc2 across prefrontal neurons in animals that never experience isolation produces no avoidance phenotype whatsoever - normal acquisition, normal latencies, normal freezing, normal retrieval - establishing that genetic vulnerability alone is insufficient. This strongly supports the gene-by- environment logic the work was designed to test ^82,83^: Tsc2 heterozygous loss becomes consequential only where stress has already recruited the cells in which it acts. It also bounds the phenotype behaviorally, since innate anxiety, stereotypy, and neutral recognition memory are all intact in SIStag.TSC mice, so what is altered is not affect or memory in general but the arbitration of defensive responding under threat - a specificity that may bear on the selective emergence of anxiety and emotional dysregulation within the broader TSC neuropsychiatric spectrum ^40,41^.

The stress-recruited population projects predominantly to the ipsilateral lateral septum, a structure positioned to translate prefrontal signals into defensive action selection through its hypothalamic and midbrain targets ^84–86^. Whether the maladaptive strategy is expressed through this projection remains untested, and pathway-specific manipulation is the natural next step.

## Conclusion

Pathological avoidance in these animals is not a failure of performance but a success that cannot be turned off. Stress recruits a prefrontal ensemble; within that ensemble, loss of Tsc2 blocks the degradative arm of proteostasis; and the resulting cells drive a response that is fast, effective, cue-locked, and blind to its own consequences. That this phenotype requires both the genetic lesion and the environmental one, and that it is reversible by restoring mTORC1 signaling within the ensemble itself, identifies a specific molecular and cellular locus at which adversity and genetic vulnerability converge to reshape defensive behavior.

## Author contributions

Conceptualization: P.S. and S.L. Methodology: S.L., D.S., O.T., M.M. and P.S. Investigation: S.L., D.S., O.T., and M.M. Funding acquisition: P.S. Supervision: P.S. Writing – original draft: P.S. and S.L. Writing – review & editing: D.S., O.T., and M.M.

## Acknowledgements

We thank all Shrestha Lab members for critical feedback and discussions. AAV5.cFos.Creert2 (AAV5.cFOS.Cre.ERT2.PEST.P2A) was a gift from Dr Karl Deisseroth (UNC viral prep Lot# AV395C); AAV.DIO.tdTomato (AAV PHP.eB CAG FLEX.tdTomato) was a gift from Edward Boyden (Addgene plasmid # 28306; http://n2t.net/addgene:28306; RRID:Addgene_28306)).

## Funding Statements

P.S. is supported by National Institute of Mental Health grant R01 MH132795, Alfred P Sloan Foundation fellowship FG-2022-94485. S.L. is supported by Scholars in Biomedical Sciences Program (T32GM148331).

## Materials and Methods

### Mice

Mice were provided with food and water ad libitum and were maintained in a 12h/12h light/dark cycle at stable temperature (78^0^F) and humidity (40-50%). The Tsc2 floxed mice^50^ were a generous gift from Michael Gambello at Emory University. Wildtype C57BI/6J mice (Stock #000664) and Ai14 Cre-reporter mice^49^ were purchased from Jackson labs. All experiments were conducted using both male and female mice aged 2 to 5 months. All procedures involving the use of animals were performed in accordance with the guidelines of the National Institutes of Health and were approved by the Institutional Animal Care and Use Committee (IACUC) of Stony Brook University.

### Drugs and chemicals

4-hydroxytamoxifen (Sigma, #H6278) was dissolved in DMSO at 40 mg/mL stock and finally diluted to 1 mg/mL in 1% Tween-80 and 0.9% saline. Rapamycin (LC Laboratories #R-5000) was diluted in DMSO at 50 mg/mL stock and finally diluted to 1 mg/mL in 5% Tween-80, 15% polyethene glycol 400 (PEG-400), and 0.9% saline. Rapamycin at 5 mg/kg was injected intraperitoneally for 3 days pre-training and before each behavior in the rescue mice cohort. Azidohomoalanine (AHA) (Click Chemistry Tools, #1066-100) was dissolved in ddH_2_O at a stock concentration of 100 mM and diluted to 1 mM in sterile saline. Stock solution of aqueous 32% paraformaldehyde (EMS, #15714) was freshly diluted to 4% in 0.1 M PBS for transcardial perfusions and post-fixation of brain slices.

### Stereotaxic surgeries and optic ferrule implant

Mice were anesthetized with a mixture of ketamine (100 mg/kg) and xylazine (10 mg/kg) in sterile saline (injected intraperitoneally). Surgeries were carried out on a Kopf stereotaxic apparatus (Model 942), equipped with a microinjection unit (Model 5000) or Nanojector UMP3TA (WPI). The viral vector carrying AAV5-cFos-CreERT2 and AAV-CAG-DIO-tdTomato (200 nl) were bilaterally injected into the mPFC [coordinates: −2.00mm anterior posterior (AP), +/-0.25mm mediolateral (ML) and −1.60mm dorsoventral (DV)] of all mice. For mice designated for fiber photometry, viral vector carrying AAV5-cFos-CreERT2 and AAV9-hSyn-DIO-GCaMP8m (250 nl) were bilaterally injected into the mPFC [coordinates: −2.00mm anterior posterior (AP), +/-0.25mm mediolateral (ML) and −1.60mm dorsoventral (DV)]. This is followed by unilateral optic ferrule [Doric 400/430, 0.66NA, 3 mm, MF1.25 (FLT)] implant in the mPFC [coordinates: −2.00, −0.25, −1.50 (0.10 mm above DV coordinate for AAV)]. For mice designated for shRaptor knockdown, viral vector carrying AAV5-cFos-CreERT2, AAV-CAG-DIO-tdTomato, and a pSICO-based U6-shRNA[Raptor] lentivirus were bilaterally injected into the mPFC [coordinates: −2.00mm anterior posterior (AP), +/-0.25mm mediolateral (ML) and −1.60mm dorsoventral (DV)]. Upon completion of intracranial injections and/or ferrule implant, the incision of the scalp was closed using a tissue adhesive (3M Vetbond) or Metabond dental cement (Parkell, # S396) for implants. Normothermia during surgery was maintained at 36.6o 431 C by resting the mouse on a covered heating pad (7 cm X 7 cm) connected to a rodent warmer console (Stoelting, # 53800M). To prevent somatic dehydration, each mouse was subcutaneously injected with 500 μl of sterile saline before surgery, and further provided with hydrogel (Clear H2O, # 70-01-5022) in the recovery cage. General monitoring and analgesia using 3 mg/kg subcutaneous Ketoprofen (Zoetis) were provided up to three days post-surgery.

### Social isolation stress and genetic access of stress-activated neurons

Mice were socially isolated by removing them from their group-housed cage and singly housed in a new cage (1 mouse per cage) for 10 days before starting behavior testing. Mice were injected intraperitoneally with 4-hydroxytamoxifen (1mg/mL), administered 10 mg/kg body weight on the first day of social isolation (6 hours following isolation) to activate gene expression. They were provided with food and water ad libitum and maintained in a 12h/12h light dark cycle at a stable temperature (78^0^F) and humidity (40 - 50 %). No environmental enrichment was provided, the cage condition was the same as the group housed cage, and handing of the mice were minimized. At the end of the 10-day isolation, the mice were subjected to behavioral experiments or euthanized to perform histology.

### Behavior

All behavior sessions were conducted during the light cycle (7:00 AM - 7:00 PM EST). Both male and female mice were included in behavior experiments. SIStag.WT and SIStag.TSC mice were randomly assigned to experimental conditions including drug and vehicle administration, and for the order of testing in any given paradigm. Behavior experiment data were collected by experimenters that were blind to the experimental conditions and genotype. Following weaning, mice were group-housed with up to three sex-matched littermates in standard cages (8 in X 13 in X 5 in) fitted with corn cob bedding and Enviro-dri enrichment. Acclimation of the mice in the behavior room lasted at least 30 min prior to testing. A white noise machine (Yogasleep Dohm; 58 dB at a distance of 6 ft from the machine) was used to blunt any environmental noise that may induce confounding anxiety. All behavioral equipment was cleaned with 30% ethanol between animals to remove olfactory cues. When multiple paradigms were performed, testing proceeded in order from least to most demanding to minimize carryover stress between assays.

#### Open field test

General locomotor activity and anxiety-like behavior were assessed in an open field test. Each mouse was placed in the center of a square open-field arena (27.31 × 27.31 × 20.32 cm) and allowed to explore freely for 15 min. Throughout each session, the animal’s spontaneous exploratory movement was recorded and tracked automatically by a computer-operated video tracking system (EthoVision XT15), which detects the animal’s body center against the arena background and reconstructs its trajectory across the session. Two parameters were extracted from the tracked trajectories. Total distance traveled-the cumulative path length of the body-center coordinate over the 15 min session-served as an index of general locomotor activity, providing a control measure to ensure that any group differences in other behavioral assays were not confounded by baseline hyperactivity or hypoactivity. The center-to-total time ratio-the proportion of session time spent in a defined central zone of the arena relative to the total session time-served as an index of anxiety-like behavior.

#### Elevated plus maze (EPM)

Anxiety-like behavior was further assessed using the elevated plus maze, a paradigm that exploits the natural conflict between a rodent’s drive to explore novel environments and its innate aversion to open, elevated, and exposed spaces. The maze was configured in a plus shape from two opposing pairs of arms - open and closed - each measuring 30 × 5 cm, radiating from a shared 5 × 5 cm central platform. The closed arms were enclosed by 17 cm high opaque walls that provide a sheltered, protected environment, whereas the open arms lacked walls entirely, leaving the animal exposed on an elevated surface. The maze was raised above the floor on a support stand and was cleaned between animals to eliminate residual olfactory cues. Each mouse was placed on the center zone, typically facing an open arm, and allowed to explore the entire maze freely for 10 min. Testing was conducted under controlled, uniform lighting in a quiet room to minimize external disturbances. Throughout the session, the animal’s spontaneous locomotion was recorded and tracked automatically by a computer-operated video tracking system (EthoVision XT15), which reconstructs the animal’s trajectory and scores arm entries and occupancy based on the position of the tracked body center relative to the defined maze zones. The duration, frequency, and latency to enter each arm were tracked.

#### Self-Grooming

Self-grooming behavior was assessed in a transparent acrylic cylinder (13 cm diameter). On the first day, each mouse was habituated to the cylinder for 10 min to reduce novelty-induced reactivity. On the second day, behavior was recorded for 10 min, with the cylinder covered by a transparent plastic lid to prevent escape while preserving an unobstructed view for video capture. Grooming behavior was subsequently scored manually from the recordings, with two measures quantified: the latency to initiate the first grooming bout and the total duration of grooming across the session.

#### Novel object recognition (NOR)

Mice were tested for novel object recognition in a 46 × 35 cm arena. During habituation (days 1–3), two identical familiar objects (green Lego blocks) were placed at the center of each arena half, and mice explored freely for 10 min/day^87^. On day 4 (long-term memory retrieval), the familiar object on each mouse’s less-preferred side (determined from habituation exploration) was replaced with a distinct novel object (a white cone), and mice explored for 5 min, with novel-object side counterbalanced across animals to control for side bias. Sessions were recorded from an overhead Logitech camera at 640 × 480, transcoded to H.264/MP4 with FFmpeg (preserving native frame rate), and tracked by markerless pose estimation using DeepLabCut ^88,89^(v3, PyTorch, ResNet-50 backbone from ImageNet-pretrained weights). Two body parts were labeled-the head (for object investigation) and body center (for locomotion)-using ∼70 k-means-clustered frames across seven representative videos, manually annotated and trained at a 0.8 training fraction with a 0.6 likelihood threshold. Training and inference ran on NVIDIA V100 GPUs of the SeaWulf HPC cluster (Stony Brook University) via SLURM, yielding per-frame x–y coordinates and likelihoods for each body part.

Behavioral metrics were quantified from tracked coordinates in custom Python software (OpenCV, NumPy, pandas). The arena boundary was delineated in a representative frame to calibrate pixel-to-centimeter conversion from known dimensions, and objects were localized by color/shape segmentation (HSV thresholding for the green familiar object; circular detection with image-moment centroid refinement for the novel object), each assigned as familiar or novel by its left–right position matched to the counterbalanced placement. A zone of interaction was defined 1 inch (2.54 cm) beyond each object’s outer edge. Investigation was scored frame-by-frame whenever the head coordinate (constrained to the nearest boundary point if outside the calibrated arena) fell within a zone, yielding three metrics per object: duration (in-zone frames ÷ frame rate), frequency (number of discrete bouts), and latency (time to first zone entry; animals never entering were excluded from latency measures for that object). Total distance traveled, computed as cumulative frame-to-frame Euclidean displacement of the body center converted to centimeters, served as an independent locomotor measure. Recognition memory was quantified by a novelty preference index for each metric, *PI = (NO − FO) / (NO + FO)*^90^, where positive values reflect preferential novel-object investigation (intact recognition memory) and values near zero indicate no preference; the index was left undefined when NO and FO summed to zero or a value was absent. Group comparisons were based on the duration-derived preference index and the latency to investigate the novel object

#### Signaled active avoidance (SAA)

##### Data acquisition

Behavioral experiments were performed using the Habitest Linc System (H01-01 Power Base and H02-08 Linc Boxes; Harvard Apparatus) controlled by Graphic State 4 software (Coulbourn Instruments), which regulated delivery of footshock, light stimuli, and auditory tones. Graphic State 4 also recorded shuttling behavior throughout each session. Freezing during SAA training and memory retrieval was quantified using FreezeFrame 4 software (Actimetrics), which captured video and automatically scored freezing responses. Two distinct environmental contexts (Contexts A and B) were used for the SAA paradigm. Context A consisted of a chamber (29 cm W × 23 cm D × 20 cm H) equipped with a white house light and metal shock grid floor, cleaned with 30% ethanol, and lacking striped wall panels. Context B was a chamber of similar dimensions but contained infrared illumination, striped wall panels, a white plexiglass platform floor, vanilla-scented paper bedding, and was cleaned with 30% isopropanol to provide distinct visual, tactile, and olfactory cues. Chambers were housed within a sound-attenuating isolation cubicle (30W × 19H × 18D; H10-24 and H10-24A, Coulbourn Instruments). Footshocks were delivered via a Precision Animal Shocker (H13-15, Coulbourn Instruments), and auditory stimuli were presented programmatically using an H12-01M speaker module coupled to an H12-07 seven-tone audio cue.

##### Context habituation

Prior to conditioning, each mouse received a single 15 min habituation session in the training context (Context A) to attenuate neophobia and reduce non-associative reactivity (e.g., stress-induced hypoactivity or freezing) that could otherwise confound acquisition of the avoidance response. Habituation without any CS or US presentation allowed mice to acclimate to the apparatus, the shuttle configuration, and the ambient sensory environment, establishing a stable behavioral baseline against which conditioned responding could be measured.

##### Pavlovian trial

On the first day of training, acquisition was initiated with a single unavoidable Pavlovian trial designed to instantiate the CS-US association before instrumental avoidance contingencies were introduced. This trial consisted of a 15 s conditioned stimulus (CS; a 5 kHz, 80 dB pure tone) that co-terminated with a 0.5 s footshock unconditioned stimulus (US; 0.2 mA) delivered through the grid floor in Context A. Because shuttling had no programmed consequence on this trial, the animal could not escape or avoid the US, ensuring that every mouse experienced the full CS-US pairing and entered the instrumental phase with an equivalent Pavlovian learning history.

##### Instrumental avoidance training

The priming trial was followed by 11 avoidance trials in Context A, delivered with a variable inter-trial interval averaging 1 min (jittered around this mean to prevent temporal prediction of trial onset). On each trial, the tone CS was presented for up to 15 s. A shuttle response (crossing from one compartment to the other) during the CS immediately terminated the tone and cancelled delivery of the scheduled US; such trials were scored as successful avoidance responses. If the animal failed to shuttle before the 15 s CS elapsed, the footshock US was delivered at CS offset, and the trial was scored as a failure. This contingency establishes a negatively reinforced instrumental response in which the CS acquires a warning-signal function and the shuttle becomes an active coping response that pre-empts the aversive outcome. Mice underwent this training protocol once per day for three consecutive days (SAA1-SAA3), permitting quantification of within- and across-session learning as the proportion of successful avoidance trials increased over training.

##### Long-term memory (LTM) retrieval tests

Recent long-term memory was probed 24 h after both the first (SAA1) and third (SAA3) training sessions, allowing assessment of avoidance memory following initial acquisition versus consolidated, multi-session learning. To dissociate the learned CS-response association from the training context and its constituent cues, LTM tests were conducted in a novel environment (Context B) that differed from Context A across every sensory modality: a white plexiglas platform floor, vanilla-scented bedding, infrared illumination (in place of the white houselight), and striped wall inserts. During each LTM test, mice were presented with five CS presentations in the absence of the US, providing a shock-free readout of conditioned responding to the tone. This design measures both the animal’s tendency to emit the learned avoidance (shuttle) response to the CS and any competing defensive freezing evoked by the tone, while the context shift ensures that responding reflects the cued association rather than contextual conditioning to Context A.

##### Response acquisition and scoring

Avoidance responses (shuttle latencies, success/failure classification, and per-trial avoidance rates) were detected and logged automatically using Graphic State 4 software, which controlled trial timing and stimulus delivery. Freezing - defined as the absence of all movement except that required for respiration - was quantified concurrently from the same sessions using FreezeFrame 5 software, providing a complementary index of passive (Pavlovian) defensive responding alongside the active avoidance measure.

### Immunohistochemistry

40 µm coronal sections of fixed brains were generated using Leica vibratome, which were subjected to immunolabeling with primary antibodies for Rat anti-mCherry (Thermofisher #M11217), Chicken anti-GFP (Abcam, #ab13970), Rabbit anti-Tsc2 (Atlas Antibodies #HPA030409), Rabbit anti-Raptor (Cell Signaling #2280S) and Rabbit anti-p62 (Abcam #ab91526). The following Alexa-Fluor conjugated secondary antibodies were used: AF568 Anti-Rat (Thermo Fisher # A-11077), AF488 Anti-Chicken (Thermo Fisher, # A32931), and AF488 Anti-Rabbit (Thermo Fisher, #A11008). Imaging data from immunohistochemistry experiments were acquired using a laser scanning microscopy (LSM) confocal microscope (Zeiss) with 20X objective lens (with 1X or 2X zoom) and z-stacks (approximately 6 optical sections with 0.563 μm step size) for three coronal sections per mouse.

### Fiber photometry

To record bulk calcium activity from the prelimbic cortex (PL), a 400 μm core optic fiber (with its associated ferrule) was chronically implanted so that the fiber tip terminated immediately dorsal to the PL region previously transduced with the genetically encoded calcium indicator GCaMP8m^54^. Fiber photometry^91^ was performed using a Tucker-Davis Technologies RZ10x integrated processor controlled by Synapse software. The RZ10x consolidates the core components of a photometry rig - excitation LEDs, their current drivers, and the photodetector (photoreceiver) - into a single real-time processor, eliminating the need for separate external light sources and lock-in amplifiers. During recording, the implanted ferrule was coupled via a patch cord to the system’s fiber-optic pathway: excitation light (465 nm) was delivered down the fiber to the PL, and the resulting emission from the calcium indicator was collected back through the same fiber and routed to the onboard photoreceiver, which converts the emitted photons into a voltage signal digitized by the processor. Synapse coordinated stimulus timing, LED modulation, and continuous data acquisition throughout each session. A separate calcium-independent (isosbestic; 405 nm) excitation channel is acquired simultaneously as a control^92,93^. Each excitation channel is sinusoidally modulated at a distinct frequency, and the processor demodulates the combined photoreceiver signal to recover the independent contributions of each channel in real time. The isosbestic channel provides a reference for signal artifacts that are not related to calcium dynamics - such as photobleaching, movement, and fluctuations in fiber coupling - and is used during analysis to correct the calcium-dependent signal. After the patch cord was attached to the implanted fiber, each mouse was placed into the SAA chamber, and photometry data were acquired continuously as the animal underwent the signaled active avoidance protocol described above. Behavioral event timestamps (e.g., CS onset/offset, shuttle responses, and US delivery) were synchronized with the photometry stream through the shared acquisition system, enabling later alignment of PL calcium activity to discrete trial events (avoidance responses, failures, and freezing epochs). Raw photometry traces were exported from Synapse and processed offline using custom-written Python code.

### Fluorescent non-canonical amino acid tagging (FUNCAT)

300 μm-thick brain slices containing medial prefrontal cortex (mPFC) were prepared in cold (4°C) carbo-oxygenated (95% O_2_, 5% CO_2_) cutting solution (110 mM sucrose, 60 mM NaCl, 3 mM KCl, 1.25 mM NaH_2_PO_4_, 28 mM NaHCO_3_, 5 mM Glucose, 0.6 mM Ascorbate, 7 mM MgCl_2_ and 0.5 mM CaCl_2_) using a VT1200S semi-automated vibratome (Leica). Slices were recovered in oxygenated artificial cerebrospinal fluid (ACSF: 125 mM NaCl, 2.5 mM KCl, 1.25 mM NaH_2_PO_4_, 25 mM NaHCO_3_, 25 mM glucose, 1 mM MgCl_2_ and 2 mM CaCl_2_) at 32°C for 2h before 1mM azidohomoalanine (AHA) was added to the ACSF solution for an additional 2.5 h. Following incubation, the mPFC areas were dissected out and submerged in 4% paraformaldehyde at 4°C overnight. The fixed micro-slices were embedded in 3% agarose, re-cut into 40 μm sections using the vibratome, and stored at −20°C in the cryosolution overnight. The mPFC sections were blocked and permeabilized in 5% IgG protease-free BSA, 3% normal goat serum, 0.3% Triton X-100 in 1X PBS at RT for 90 min with agitation. The brain sections were subjected to click chemistry using Cell Reaction kit (Thermo Fisher #C10269) with 25 μM Alkyne Alexa fluor 647 (Invitrogen, #A10278), CuSO_4_ and additive overnight at 4°C. The following day, the sections were washed 3 times with 1X PBS for 10 min per wash. Sections were blocked with 1% NGS in 1X PBS for 1h followed by incubation with the primary antibody: anti-mCherry (Thermofisher #M11217) overnight at 4°C for standard immunohistochemistry.

### Imaging

Immunolabeled tissue was imaged on a Zeiss LSM 800 laser scanning confocal microscope using a 20× objective (1× or 2× zoom), acquiring z-stacks of approximately six optical sections at a 0.563 μm step size across three coronal sections per mouse to sample the region of interest and reduce section-to-section variability. Acquisition settings (laser power, gain, offset) were held constant across all sections and animals within an experiment to permit valid quantitative comparison. Image stacks were imported into ImageJ (FIJI) using the Bio-Formats plugin, and maximum-intensity projections were generated from each stack.

For cell-based immunofluorescence analysis, individual cells were manually outlined as regions of interest (ROIs) using the ImageJ drawing tools, and mean fluorescence intensity was measured within each ROI. Background fluorescence was sampled from five cell-free ROIs of comparable area in the same channel and averaged; this value was subtracted from the mean intensity of each cell ROI. Background-corrected intensities were then normalized to the mean value of the wild-type (WT) control within each experiment.

Cell size was quantified by manually outlining the perimeter of each labeled cell using the FIJI drawing tools and measuring the enclosed area (µm²). To assess dendritic morphology, dendrites from each labeled cell were manually reconstructed using the Simple Neurite Tracer plugin in FIJI. All visible primary and higher-order dendritic branches originating from the cell body were traced throughout the image stack, and the cumulative length of all traced dendrites was calculated to obtain a single total dendritic length measurement for each cell. This approach provided a quantitative measure of dendritic arborization that was used for comparisons across experimental groups. Tracing was performed using identical image display settings and analysis parameters across all samples to ensure consistency. Where dendrites extended beyond the image boundaries or could not be unambiguously resolved, only the visible portion of the dendrite was included in the analysis.

To quantify labeled axonal projection density in the lateral septum (LS), where signal is distributed across neuropil rather than discrete stomata, an integrated fluorescence approach was used. From the maximum-intensity projections, an ROI encompassing the LS was manually delineated, and integrated density (ROI area × mean gray value) was measured with the ImageJ tools. To correct for non-specific background, mean intensity was sampled from the same region in Ai14 control tissue devoid of specific signal, and background-corrected integrated fluorescence was calculated as integrated density − (ROI area × mean background intensity). Corrected values were averaged across sampled sections to yield a single projection-signal measure per animal for comparison across experimental groups.

### Data Analysis

#### Fiber Photometry Data Analysis

To preprocess and analyze fiber photometry recordings from signaled active avoidance (SAA) experiments, we implemented a multi-step artifact removal and signal normalization pipeline using custom Python software. Raw TDT data were imported using the Python library tdt via a custom data loading function, which reads the .tev and .tsq files automatically generated in the specified tank directory. To capture the relevant behavioral epochs while excluding extraneous signals, time windows were tailored to each session type: for SAA training sessions containing eleven CS presentations, signals were extracted beginning 120 seconds prior to the second CS onset (excluding the first habituation trial) through the end of the session; for long-term memory (LTM) retrieval sessions, signals were extracted beginning 120 seconds prior to the first CS onset. This approach ensured that all CS-evoked neural responses were captured within the analysis window while minimizing the inclusion of pre-task baseline periods unrelated to the behavioral paradigm.

Following data import, a second-order Butterworth low-pass filter with a 4 Hz cutoff frequency was applied to both the calcium-dependent signal channel (465 nm, GCaMP8m) and the isosbestic control channel (405 nm) to suppress high-frequency noise and artifacts that can obscure the slower, biologically meaningful fluorescence dynamics associated with neural calcium transients. Because fiber photometry signals are susceptible to gradual fluorescence decay over the course of a recording session, photobleaching correction was performed by fitting a double exponential decay function to each channel independently and subtracting the resulting fit from the filtered traces^94,95^. This step effectively removed the slow baseline drift caused by progressive fluorophore bleaching, preserving the fidelity of stimulus-evoked responses. To address motion artifacts arising from animal movement during the behavioral task, we applied a linear regression-based motion correction strategy. The linear relationship between the isosbestic control channel (405 nm) and the calcium-dependent signal channel (465 nm) was modeled by ordinary least squares regression, and the predicted signal derived from the control channel was subtracted from the photobleaching-corrected signal trace. This widely adopted approach minimizes non-neural, movement-induced fluorescence fluctuations that are present equally in both channels. Following motion correction, we computed ΔF/F, defined as the percentage change in fluorescence relative to a fitted baseline fluorescence value: ΔF/F = (F − F_baseline) / F_baseline × 100, where F represents the motion-corrected fluorescence signal and F_baseline is the photobleaching fit of the signal channel.

To enable comparisons across sessions and animals, ΔF/F values were converted to z-scores using the transformation z = (x − μ) / σ, where x is each ΔF/F value, μ is the session mean, and σ is the session standard deviation of the ΔF/F distribution within the analyzed time window. A positive z-score reflects calcium activity above the session mean, whereas a negative z-score indicates suppressed fluorescence relative to baseline. To smooth the normalized signal while retaining key features such as stimulus-evoked peaks and transients, we applied a Savitzky–Golay filter^96^ with a 1001-frame sliding window and polynomial order 3. This filter fits local polynomials over a sliding window, generating convolution coefficients that minimize squared error and reduce frame-to-frame variability without distorting the underlying signal dynamics. This smoothing step improves both visualization and downstream quantitative analyses while preserving biologically relevant temporal features.

#### Trial Classification and Peri-Event Time Histograms

Individual CS presentations were classified as either failed or successful trials based on CS duration. Failed trials were defined as those in which the CS persisted for the full duration (≥ 14.9 s), indicating that the animal did not shuttle to the opposite chamber and received the unconditioned stimulus (US, foot shock). Successful trials were defined as those in which the animal performed an active avoidance response (AR) prior to US delivery, resulting in early CS termination (< 14.9 s); the AR time was operationally defined as the moment of CS offset. Peri-event time histograms (PETHs) were computed separately for failed and successful trial types, aligned to CS onset. For failed trials, z-scored fluorescence traces were extracted over a fixed 45-second window (15 seconds before CS onset to 30 seconds after CS onset), capturing the pre-CS baseline, the full 15-second CS period, and the post-CS epoch following US delivery. For successful trials, because CS duration varied across trials and animals depending on when the avoidance response occurred, a time-warping procedure was applied to enable meaningful across-trial averaging. Specifically, the CS period of each trial’s fluorescence trace was linearly interpolated to a common target duration equal to the group-level median CS duration, computed by pooling all successful trials across SAA sessions (SAA1, SAA2, SAA3) or LTM sessions (LTM1, LTM2) within each experimental group. The pre-CS (15 seconds) and post-AR (15 seconds) epochs flanking the warped CS period retained their original temporal resolution. This time-warping approach^55^ aligns the avoidance response across trials at a common reference point, allowing the neural dynamics preceding and following the AR to be compared without temporal misalignment introduced by variable CS durations. For each animal, mean ± SEM PETH traces were computed across all trials of each type within a session and plotted to visualize session-level neural responses. For group-level analyses, individual animal PETHs were averaged within predefined experimental groups (SIStag.GCaMP8m and SIStag.TSC2cHET.GCaMP8m) to enable comparisons across genotypes.

##### Quantification of Neural Responses

To quantify stimulus-evoked neural responses for statistical comparisons, two complementary metrics were computed from the z-scored fluorescence traces. First, the peak calcium transient (Ymax) was defined as the maximum z-score value within each epoch of interest for each individual trial, and per-animal means were computed by averaging across trials within each session. Second, the area under the curve (AUC) was calculated using the trapezoidal integration method over defined epoch windows.

For failed trials, three 15-second epochs were defined: Pre-CS (−15 to 0 s relative to CS onset), CS (0 to +15 s), and Post-CS (+15 to +30 s). For successful trials, to enable direct statistical comparison across epochs, epoch durations were matched to the group-level median CS duration. Specifically, the Pre-CS epoch spanned from −group_median to CS onset, the CS epoch spanned the time-warped CS period (0 to group_median), and the Post-CS epoch spanned an equivalent matched window immediately following the AR (AR to AR + group_median). In addition, a 15-second post-AR AUC was computed and subdivided into three consecutive 5-second sub-epochs ([AR to AR+5s], [AR+5s to AR+10s], [AR+10s to AR+15s]) to characterize the temporal dynamics of neural activity following the avoidance response. Group-level median CS durations used for epoch matching were identical to those used for the time-warping procedure described above, ensuring consistency between visualization and quantification.

#### Behavioral data analysis

To facilitate efficient and reproducible analysis of avoidance conditioning experiments, we developed a fully automated, Python-based computational pipeline for data extraction, behavioral metric computation, and visualization. The workflow integrates timestamped event logs from FreezeFrame (freezing during Pavlovian threat conditioning and differential signaled active avoidance) and Graphic State (threat-predictive conditioned stimulus [CS], and unconditioned stimulus [US] - evoked responses) to generate structured, analysis-ready datasets. Custom modules (freezeframe.py, graphicstate.py) implement preprocessing steps including column name standardization, epoch segmentation (pre-CS, CS for freezing behaviors, and non-CS, CS for shuttling responses), metadata integration, and handling of missing data. Task-specific extensions (saa_gs.py) were generated to classify avoidance trial outcomes (avoidance, escape, failure), compute spontaneous and CS evoked shuttling, and calculate latency to avoid using precisely parsed CS onset/offset timestamps. On the other hand, saa_ff.py was generated to quantify freezing percentages across defined epochs (pre-CS, CS and postCS). The pipeline uses an object-oriented modular design built around the FreezeFrame and GraphicState base classes, supporting reusability across experimental paradigms. Automated batch processing was implemented for high-throughput analysis of multiple datasets with minimal manual input. Comprehensive logging and error handling (via logging.info(), logging.warning(), try-except blocks) were also utilized to ensure data integrity and traceability. Dedicated plotting modules (fr_pav_heatmaps.py) were developed to generate standardized visualizations of freezing dynamics, allowing rapid and reproducible figure generation. Together, this custom pipeline provided a scalable and transparent framework for behavioral data analysis in differential signaled active avoidance paradigm.

### Statistics

Statistical analyses were performed using GraphPad Prism 8 (GraphPad software). Data are expressed as mean +/- SEM. Data from two groups were compared using two-tailed Unpaired Student’s t test. Multiple group comparisons were conducted using one-way ANOVA, two-way ANOVA, or RM Two-way ANOVA with post hoc tests as described in the appropriate figure legend. Statistical analysis was performed with an α level of 0.05. P values <0.05 were considered significant.

## Notes

### Competing Interest Statement

The authors have declared no competing interest.

