## Supplementary Figures for "TSC/mTORC1-dependent proteostasis in stress-recruited prefrontal ensembles gates flexible avoidance"

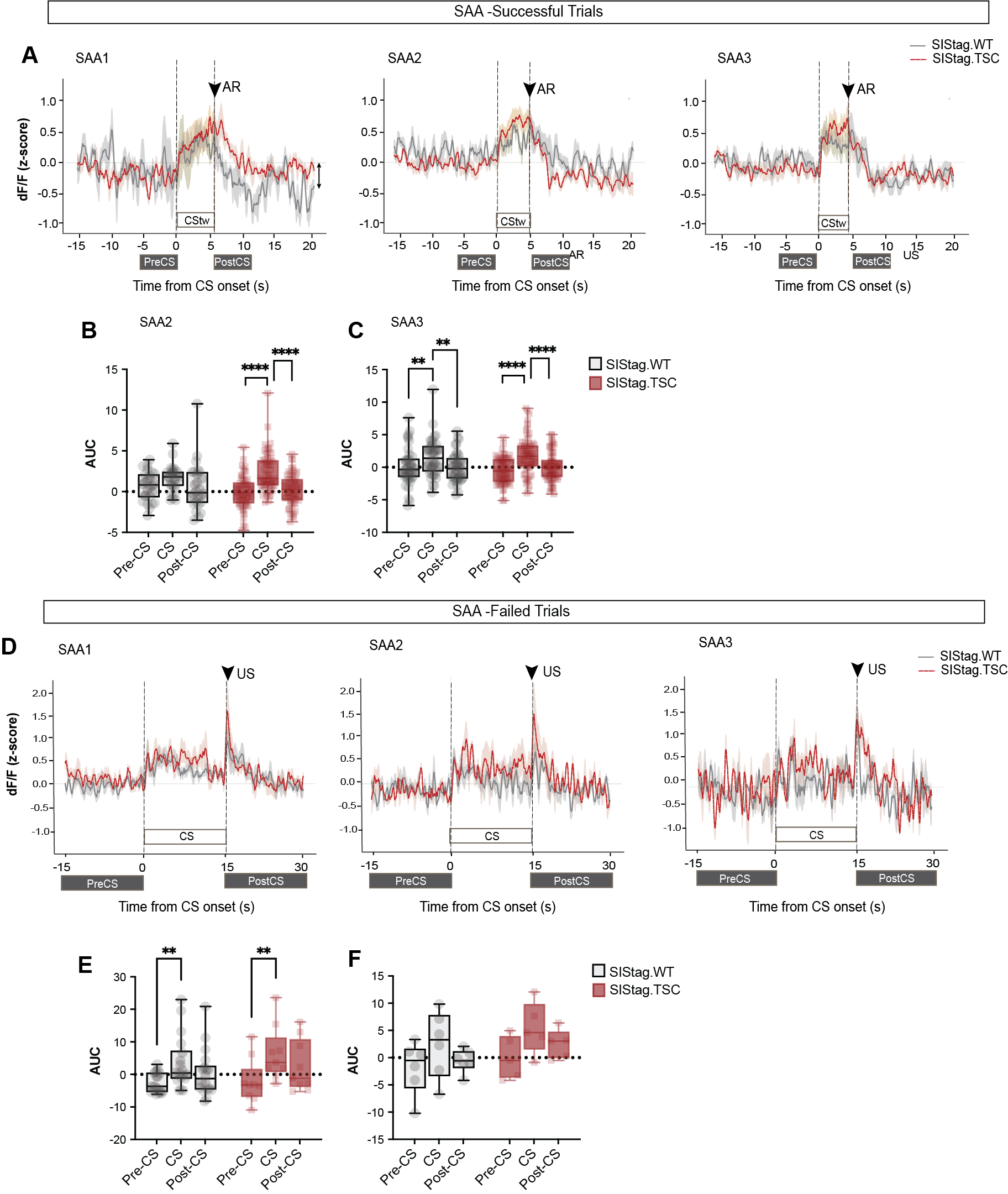

**Supplementary Figure 1. A)** Overlay of peri-event time histograms for successful trials for SIStag.WT (grey) and SIStag.TSC (red) mice during SAA1, SAA2 and SAA3. CS time-warped is marked as CStw, and PreCS as well as PostCS epochs matching the CStw duration are marked. **B)** While SIStag.WT mice do not have robust neural response to CS during SAA2, SIStag.TSC mice have elevated response to CS compared to PreCS and PostCS epochs. **C)** During SAA3, both SIStag.WT and SIStag.TSC mice have elevated responses to CS compared to PreCS and PostCS. **D)** Overlay of peri-event time histograms for failed trials for SIStag.WT (grey) and SIStag.TSC (red) mice during SAA1, SAA2 and SAA3. **E)** Both SIStag.WT and SIStag.TSC mice have elevated responses to CS presentation compared to PreCS during SAA2. **F)** The number of failed trials reduces drastically in SAA3 for both SIStag.WT and SIStag.TSC mice, and hence no significant increase in neural response is observed during CS presentation for both SIStag.WT and SIStag.TSC mice. **Stats**

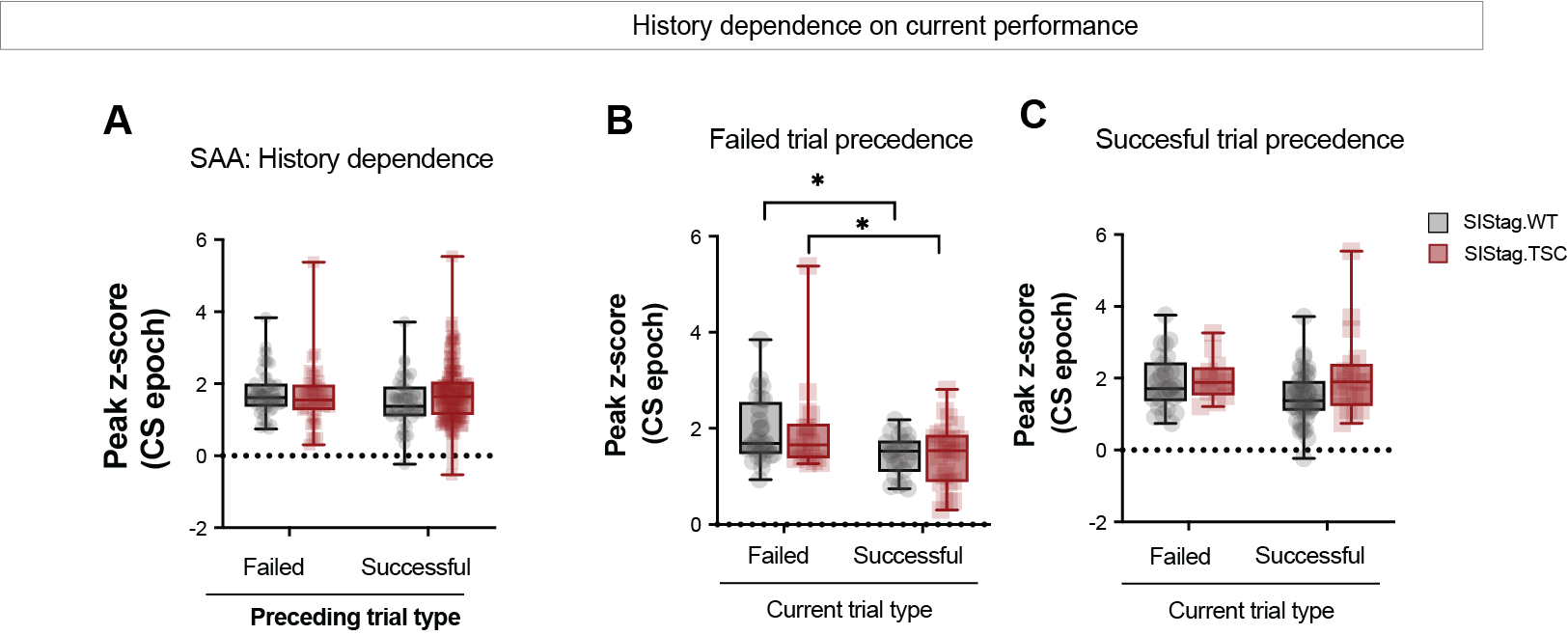

**Supplementary Figure 2. A)** History dependence analysis for neural responses of SIStag.WT and SIStag.TSC mice during SAA shows no difference between groups when the preceding trial type is failed or successful. **B)** Both SIStag.WT and SIStag.TSC mice show a decrease in neural response (Peak z-score) to CS when a successful trial is preceded by a failed trial, compared to the case when a failed trial is preceded by a failed trial. **C)** No difference is observed between current trial types when the trials are preceded by a successful trial. **Stats**

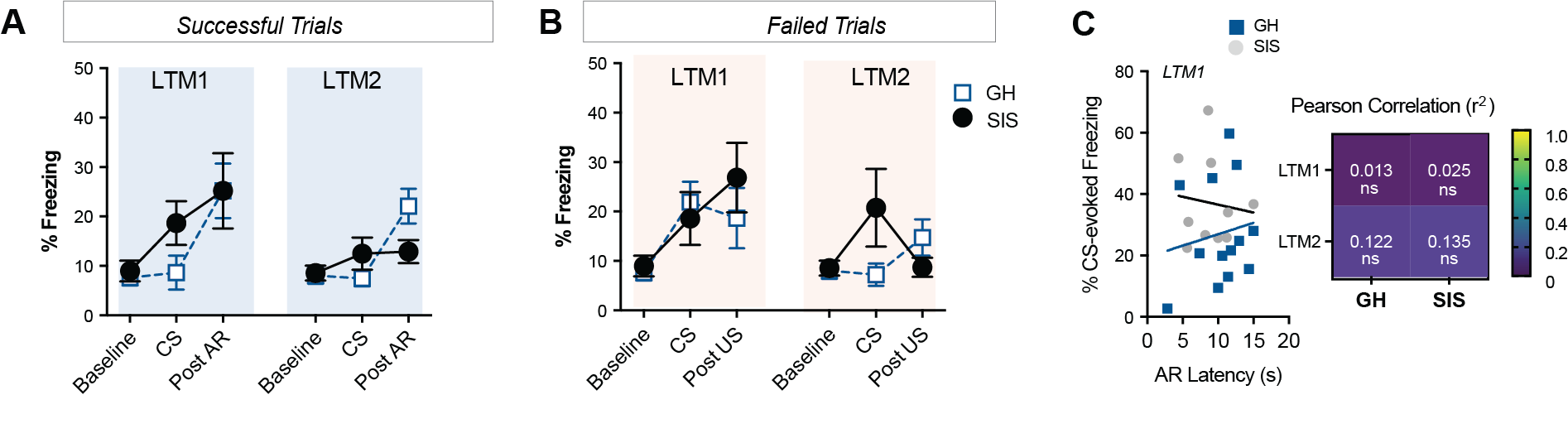

**Supplementary Figure 3. A)** In successful trials, freezing response is consistently low for both grouphoused (GH) and socially isolated (SIS) mice at baseline, CS epoch and post AR epoch for both LTM1 and LTM2 tests. **B)** In failed trials, freezing response is similarly low for both groups at baseline, CS and post US epochs for LTM1 as well as LTM2. **C)** Lack of correlation between CS-evoked freezing response and avoidance latency in GH and SIS mice. **Stats**

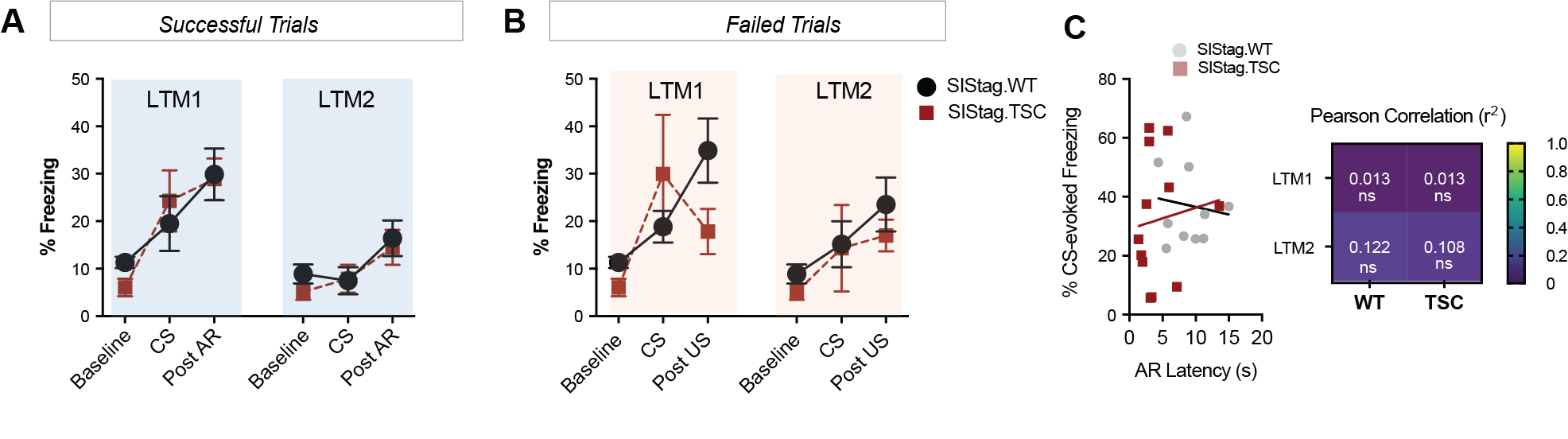

**Supplementary Figure 4. A)** In successful trials, freezing response is consistently low for both SIStag.WT and SIStag.TSC mice at baseline, CS epoch and post AR epoch for both LTM1 and LTM2 tests. **B)** In failed trials, freezing response is similarly low for both groups at baseline, CS and post US epochs for LTM1 as well as LTM2. **C)** Lack of correlation between CS-evoked freezing response and avoidance latency in SIStag.WT and SIStag.TSC mice. **Stats**

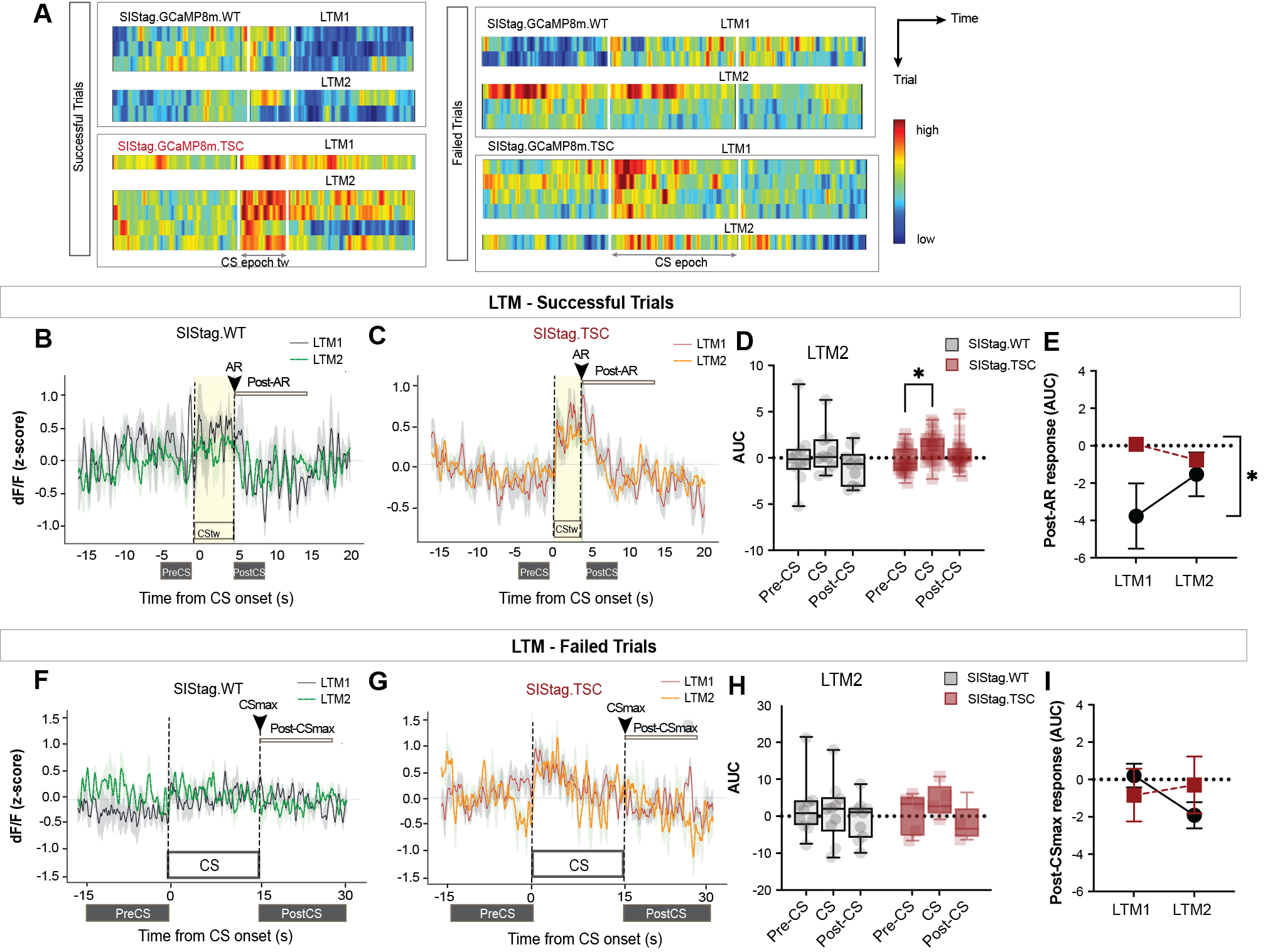

**Supplementary Figure 5. A)** Heatmaps showing intensity of neural responses in stress-activated neurons during successful and failed trials in SIStag.WT and SIStag.TSC mice expressing ensemble-specific GCaMP8m. **B & C)** PETH showing CS and avoidance evoked changes in neural responses in SIStag.WT and SIStag.TSC mice. **D)** During LTM2, only SIStag.TSC mice have elevated neural response during CS presentation compared to PreCS. **E)** Post-AR response during 10s following avoidance is significantly different for SIStag.WT and SIStag.TSC groups during LTM1 but converges during LTM2. **F & G)** PETH showing CS and postCSmax evoked changes in neural responses in SIStag.WT and SIStag.TSC mice. **H)** During LTM2, neither group shows any elevation in neural response to the CS. **E)** Post-CSmax response (AUC) is low and unchanged for both groups during LTM1 and LTM2. **Stats**

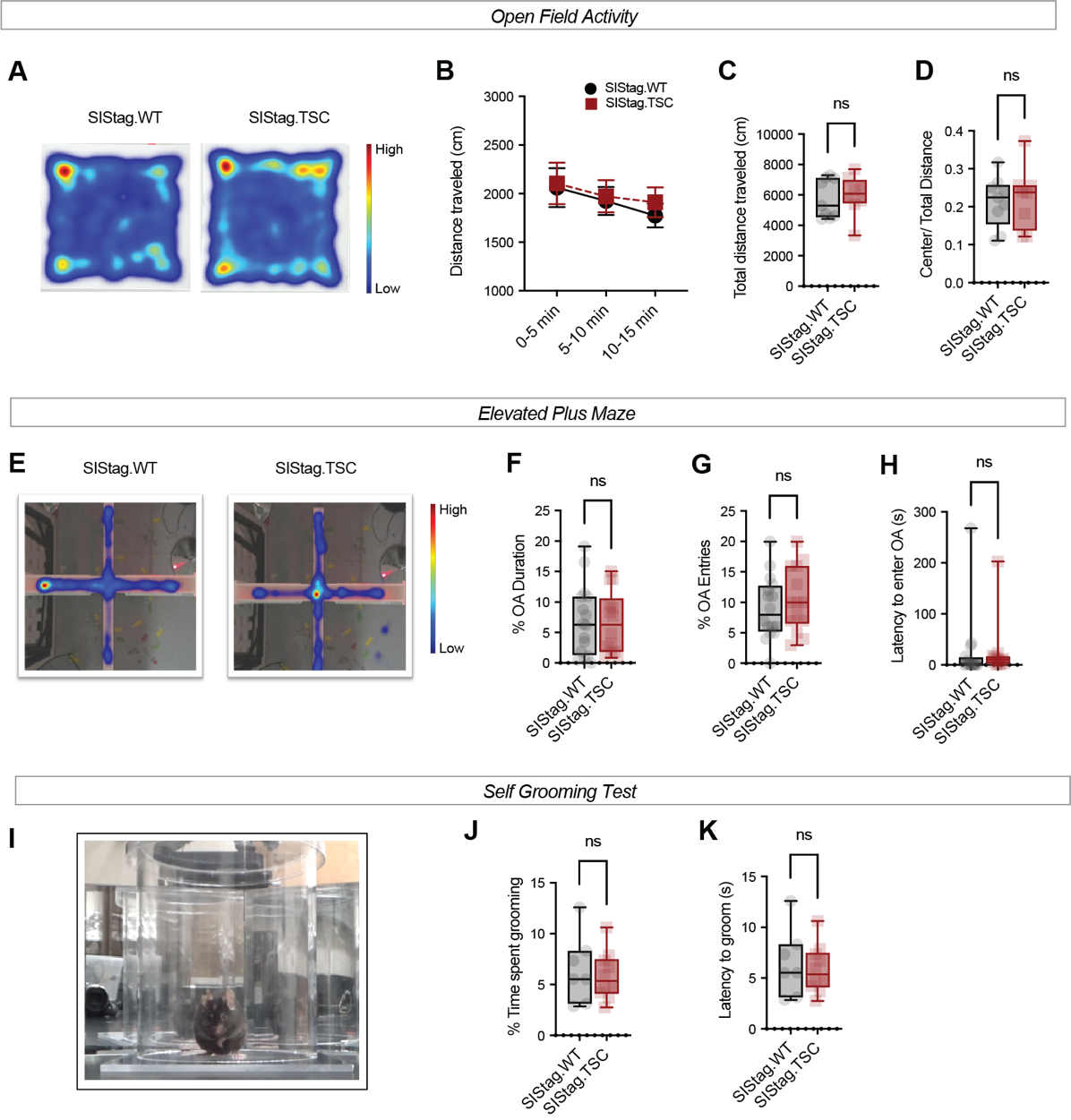

**Supplementary Figure 6. A)** Heatmap showing preferred location and spontaneous exploration of SIStag.WT and SIStag.TSC mice in the open field test. **B)** Distance traveled across time was comparable for SIStag.WT and SIStag.TSC mice, although acclimation to the arena was minimal. **C)** Total distance traveled by SIStag.WT and SIStag.TSC mice was comparable. **D)** The ratio of center to total distance (thigmotaxis) was comparable between the groups. **E)** Heatmap showing exploration of the open and closed arms in an elevated plus maze test. **F)** Percent duration in the open arm (OA) was comparable for both SIStag.WT and SIStag.TSC mice. **G)** Percent open arm entries were comparable between the groups. **H)** Latency to enter open arm was similarly comparable for the two groups. **I)** Image showing the self grooming test. **J)** Percent time spent grooming was equivalent for two groups. **K)** Latency to groom was similar for both groups. **Stats**

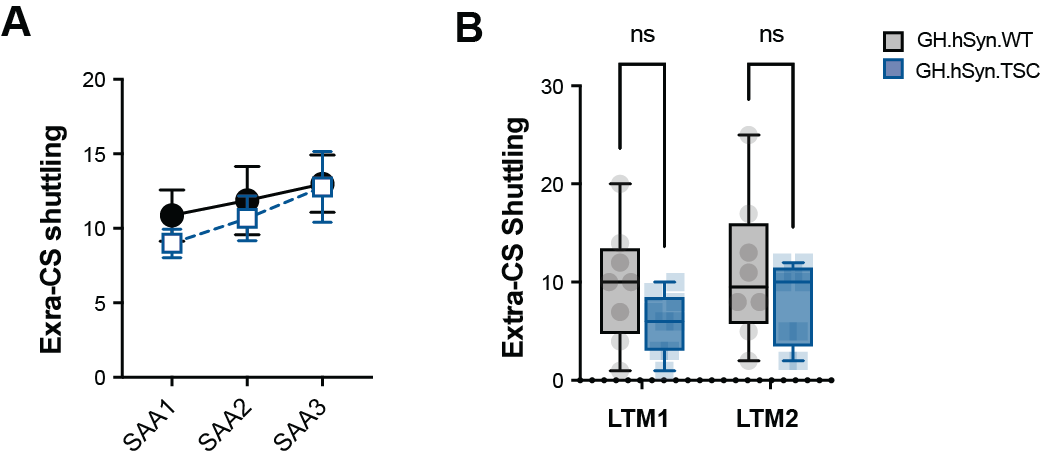

**Supplementary Figure 7.** A) Extra-CS shuttling during SAA trials (SAA1, SAA2, SAA3) for grouphoused hSyn.WT and hSyn.TSC mice showing stable responding across training. B) Extra-CS shuttling remains unchanged between the groups during LTM1 and LTM2. **Stats**
